# Fine mapping of PTSD GWAS reveals a role for amygdala *Foxp2* in regulation of fear and threat responses

**DOI:** 10.64898/2026.06.22.733764

**Authors:** O Ponomareva, LT Seabrook, M Maya-Martinez, C Klengel, V Balakundi, S Kini, E Catt, J Zion, R Shah, P Martinez, E Hernandez, Z Beatty, MS Millet, Q Flanagan-Burt, AA Lussier, CM Nievergelt, AX Maihofer, K Koenen, PTSD Working Group of Psychiatric Genomics Consortium, PsychENCODE PTSD BrainOmics Project, JE Kleinman, J Suh, WA Carlezon, NP Daskalakis, KJ Ressler

**Author notes:** Corresponding author: Kerry J. Ressler, McLean Hospital, 115 Mill St, Belmont MA 02478.

## Abstract

Post-traumatic Stress Disorder (PTSD) is a debilitating psychiatric condition caused by severe trauma exposure and characterized by ongoing dysregulation of fear processing, hyperarousal, and amygdala activation, but with limited effective treatments. Recent large-scale genome-wide association studies (GWAS) of PTSD have identified the transcription factor FOXP2 as a highly-significant, top putative risk gene. Both fine-mapping of the PTSD GWAS with amygdala-specific expression quantitative trait loci (eQTL) data, and transcriptome-wide association analyses, show that altered expression of *FOXP2* is associated with increased PTSD risk. In vertebrates, *FOXP2* mRNA is most densely expressed in the intercalated cells (ITCs) of the amygdala. ITC neurons receive excitatory input from external regulatory and sensory brain regions, as well as the basolateral amygdala, and send inhibitory projections to the central amygdala, which regulates downstream fear responses. While ITCs are critical for conditioned fear acquisition and extinction, the role of the *FOXP2* gene in modulating fear-related behaviors remains unknown. Here, we used complementary bioinformatic, molecular, circuit, behavioral, and electrophysiological approaches to characterize the function of mouse (*Foxp2*) and human (*FOXP2*) orthologs in amygdala-mediated fear learning.

To assess *Foxp2* function *in vivo*, we first used shRNA-mediated knockdown (KD) of *Foxp2* in ITC neurons of adult mice. Targeted *Foxp2* KD robustly and significantly reduced freezing (threat/fear expression) during and after auditory fear conditioning. Whole-cell recordings from individual ITC neurons revealed that *Foxp2* KD increased their intrinsic membrane excitability and action potential frequency, consistent with hypothesized enhanced inhibitory output to the central amygdala and thus reduced fear expression. This hyperexcitability was associated with reduced potassium channel conductance. Bulk RNA sequencing (RNA-seq) of mouse amygdala after ITC *Foxp2* KD confirmed decreased potassium channel transcription and revealed broader Foxp2-dependent regulation of multiple genes implicated in fear learning, including *Wnt*, *Crh* (which encodes corticotropin-releasing hormone), and neurokinin signaling pathways.

Consistent with these findings, bulk RNA-seq of medial amygdala postmortem tissue from humans with PTSD versus neurotypical controls showed decreased potassium channel transcription in samples with low *FOXP2* expression. Downstream transcriptional changes following *Foxp2* KD in the mouse amygdala also showed marked enrichment of genes identified in PTSD risk loci from the largest PTSD GWAS to date. Specifically, downregulated genes were enriched for mouse orthologs of Tier 1 PTSD GWAS risk genes. This enrichment appears to reflect subcortical *Foxp2* signaling within the amygdala, driven predominantly by decreased expression of genes lacking promoter-anchored chromatin loops. This finding suggests that Foxp2 may directly bind regulatory elements of multiple top PTSD risk genes, acting as a key regulatory node for fear-related pathways in the amygdala.

Collectively, our findings establish FOXP2 as a central transcriptional regulator of fear-related gene networks in the amygdala and potential regulatory hub for PTSD genetic risk.

## Introduction

Post-traumatic stress disorder (PTSD) is a debilitating mental illness associated with increased suicide risk, functional morbidity, and comorbidity with other psychiatric and medical conditions. PTSD results from life-threatening trauma with subsequent symptoms of avoidance, depressed mood, sleep disturbances, and intrusive thoughts and feelings related to the trauma. Decades of research have implicated the amygdala as a critical node underlying dysregulated fear, threat, hyperarousal, and decreased sense of safety in patients with PTSD.^1–7^ Recent advances in large scale genomic studies, including Genome-Wide Association Studies (GWAS) have identified *FOXP2* as the top gene putatively implicated in PTSD, with association of the most robust SNP at *p* < 5.0×10^-20^ (**Figure 1A**).^8^ Transcriptome-Wide Association Studies (TWAS) further associated downregulation of *FOXP2* with PTSD risk.^8,9^

**Figure 1.**
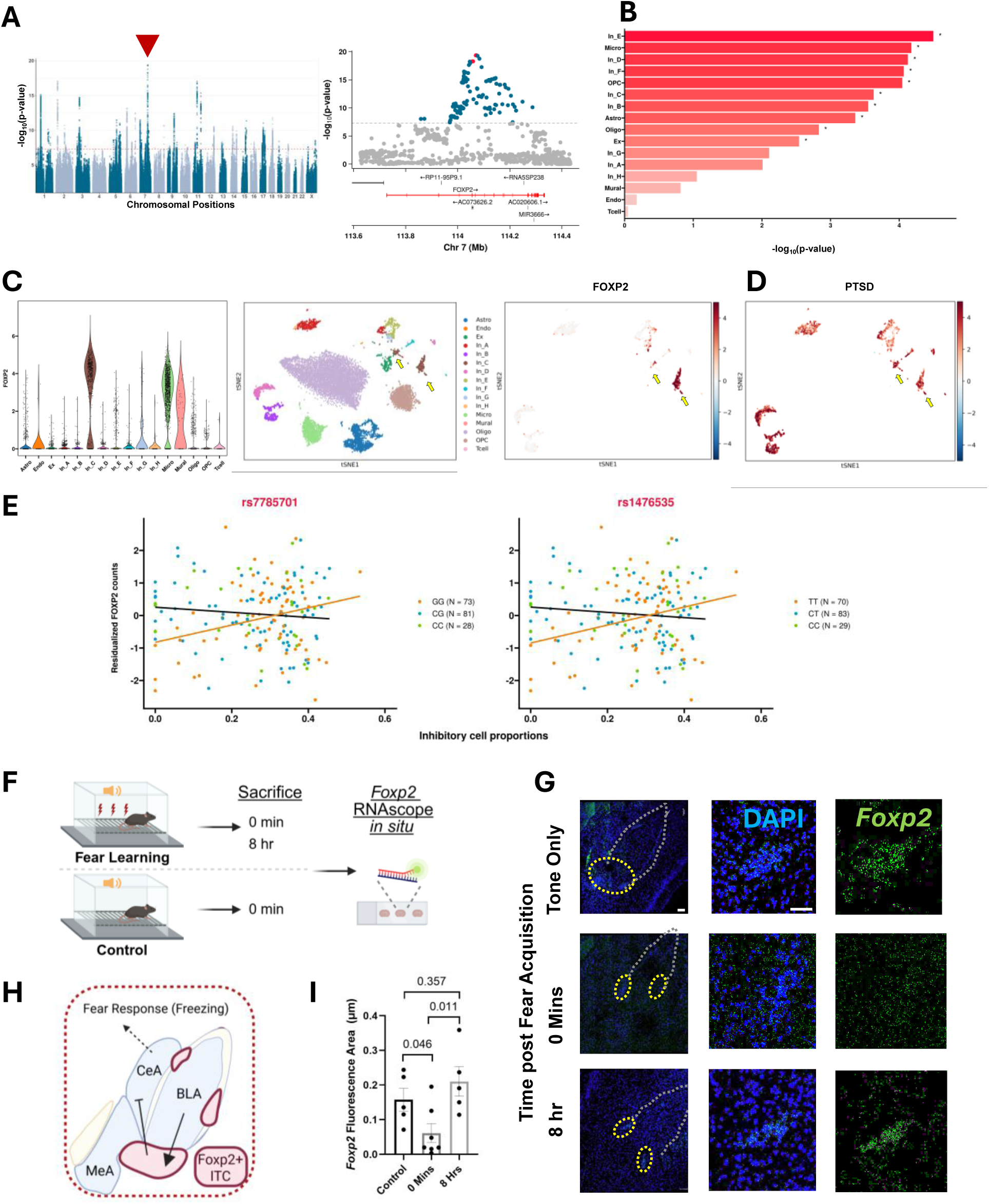
Fine Mapping of *FOXP2* locus in human PTSD GWAS studies and *Foxp2* transcriptional regulation after fear learning (i.e. threat conditioning) in mice. Fine mapping of the FOXP2 locus on chromosome 7 (red arrowhead; GWAS-significant loci associated with PTSD in European Ancestry from PGC-Freeze 3, modified from Nievergelt et al.^8^ PTSD risk-associated SNPs are highlighted in red) (A), followed by MAGMA cell type–specific enrichment analysis of PTSD GWAS risk genes (B). *FOXP2* expression levels across cell types in (C – left), along with single-cell t-SNE clustering of amygdala cells (C - middle), and distribution of *FOXP2* expression in inhibitory amygdala cells (C – right). Inhibitory neuronal population C is highlighted with yellow arrows. Disease relevance scores calculated with scDRS using PTSD GWAS gene-based associations and only the inhibitory neuronal populations in (D). (E) Residualized *FOXP2* expression is plotted against the proportion of inhibitory neurons for two SNPs within the PTSD credible set of the FOXP2 *locus* (also highlighted in red in panel A). Orange, blue, and green indicate homozygous condition for the PTSD risk allele, and heterozygous and homozygous conditions for the non-risk allele, respectively. The number of individuals (n) are reported in the legend. Linear regression lines are shown in orange for individuals homozygous for the risk allele and in black for all other individuals (heterozygous and homozygous for the non-risk alleles). We next evaluated transcriptional regulation of mouse *Foxp2* mRNA in the intercalated cells of the mouse amygdala (F). Within the amygdala, Foxp2-containing ITCs receive excitatory inputs from the BLA and send inhibitory outputs to the CeA (H). We find dynamic regulation of *Foxp2* transcription after fear learning, with *Foxp2* mRNA decreased immediately after fear learning in the ITCs of male C57Bl/6 mice (10 weeks old) (G,I). Foxp2 mRNA is labeled with RNAScope in control mice (n=5), at 0 mins (n=7) and 8 hrs (n=5) after fear learning. T-test, *p<0.05 (G). Error bars represent Standard Error of the Mean. NS, not significant. Yellow ovals represent ITC clusters. Abbreviations: Astro, Astrocytes; Oligo, Oligodendrocytes; Endo, stromal Endothelial cells; Excit, Excitatory neurons; Inhib, Inhibitory neurons; Micro, Microglia; Mural, Mural stromal cells; OPC, Oligodendrocyte progenitor cell. Excitatory and Inhibitory cell sybtypes (ie Excit_A, Excit_B, etc) represent different functional classes of neurons, marked by distinct transcription of top markers.^66^

Beyond PTSD, *FOXP2* appears to be associated with a number of transdiagnostic syndromes associated with amygdala dysfunction and emotion dysregulation. Originally characterized for its role in development, *FOXP2,* expressed robustly in the amygdala, was the first gene linked to a developmental speech and language disorder.^10^ Single nucleotide polymorphisms (SNPs) in *FOXP2* have been consistently implicated in GWAS for PTSD,^8,11^ ADHD,^12^ risk-taking behaviors,^13^ suicide risk,^14^ antisocial behavior,^15^ substance use disorders,^16,17^ self-reported childhood maltreatment,^18^ and in homozygous haplotype mapping in autism spectrum disorders.^19^ These findings suggest its mechanistic involvement in multiple psychiatric disorders, as well as in learning, memory, and reward processing. Rare heterozygous mutations in *FOXP2* lead to language disruption and an autism-spectrum phenotype, while homozygous inactivating *FOXP2* mutations are not viable into adulthood.^10,20,21^

On a molecular level, FOXP2 is a transcription factor in the forkhead homeobox family and, together with its binding partner FOXP1, is one of the most highly conserved genes in mammals.^22,23^ Beyond its role in language learning, FOXP2 is implicated in transcriptional regulation during neurodevelopment, neurite outgrowth, synapse formation, and cancer.^10,24–27^ Targeted mechanistic studies have shown that Foxp2 regulates dendrite length,^28,29^ retinoic acid signaling,^30^ *CNTNAP2* gene (a member of the neurexin family),^31,32^ SRPX2/uPAR complex (through which Foxp2 regulates synapse formation),^33,34^ *MET (*a gene implicated in autism),^35^ *DISC1* (a gene implicated in schizophrenia),^36^ *VLDLR (*a receptor for RELN),^32,37^ and the Wnt pathway.^38^ Interestingly, Wnt signaling is also involved in upstream regulation of the FOXP2/Foxp2 promoter,^39^ along with regulation by CREB and glucocorticoid signaling^40^ – pathways, which we and others have shown are implicated in fear learning in the amygdala.^41–43^ To date, five ChIP-chip and ChIP-seq studies have tested downstream transcriptional regulation by FOXP2/Foxp2, spanning human fetal tissue,^44^ human neuroblastoma cells,^26^ embryonic mouse brain tissue,^24^ neuroectodermal tumor cells and neuroblastoma cells,^45^ and male and female zebrafinch brain tissue.^46^ No identified targets were common to all studies, highlighting the importance of tissue-specificity and cellular environment on FOXP2/Foxp2 activity.

In the adult mammalian brain, *Foxp2* is expressed widely, though sparsely, in the cortex, striatum, thalamus, brainstem, cerebellum, but also very densely within a subset of amygdala cells.^47–52^ In the amygdala, *Foxp2* is strongly expressed in the intercalated cells (ITCs), a group of evolutionarily conserved cells that are also present in humans.^53–55^ GABAergic ITC neurons receive glutamatergic excitatory input from the basolateral amygdala (BLA) and other regulatory and sensory brain regions, and provide feedforward inhibition to the central amygdala (CeA), which mediates the subsequent physical response of conditioned fear via projections to the brainstem and hypothalamus.^56,57^ ITCs are also activated by stimulation of the infralimbic division (IL) of the medial prefrontal cortex (mPFC), a region critically involved in fear extinction.^58–60^ In addition to their documented role in fear extinction, more recent work suggests that a subset of ITCs is involved in the expression of conditioned fear response and feedback inhibition to the BLA.^61,62^ To date, *Foxp2* has been used as a common marker of ITCs in rodents and primates to probe the function of ITC-regulated circuits; however its molecular function in these cells is unknown.^61,63,64^ Notably, Foxp2 is a conserved transcriptional regulator in the ITCs,^55^ and thus serves as a likely hub gene for controlling molecular pathways involved in fear learning acquisition, expression, and extinction. Despite increasing genomic evidence of FOXP2’s role in psychiatric disease associated with fear/threat learning and stress responses, its role in the regulation of emotional memory processes in the amygdala – a critical brain region involved in a wide array of behaviors and psychiatric disorders, including PTSD - remains unknown.

We present the first evidence that *Foxp2* gene in the ITCs regulates fear expression and downstream transcription in the amygdala. Combining molecular, behavioral, and electrophysiological recordings, along with human genomic data analyses, our data highlight *Foxp2* in amygdala ITCs as a possible hub gene underlying fear, threat, and stress related disorders.

## Materials and Methods

### Fine Mapping of human FOXP2 gene and scDRS analysis

Using summary statistics from the Nievergelt et al. PTSD GWAS^8^, we performed gene mapping with FUMA^65^ to obtain gene-level associations. Single-cell RNA-seq data, generated by Tran et al^66^, from amygdala were used to identify cell-type-specific marker genes using the findMarkers function from the scran^67^ package. For the 19 reported cell types, astrocytes and excitatory neuron subtypes were collapsed into broader astrocyte and excitatory neuron categories, respectively, resulting in a final set of 16 cell types. To assess whether genetic associations were enriched within specific cell types, we performed gene set enrichment analysis using MAGMA^68^ with marker genes for each of the 16 cell types, selecting them based on a false discovery rate (FDR)-adjusted p-value < 0.05 threshold.

We applied single cell Disease Relevance Score (scDRS)^69^ to compute per-cell disease relevance scores based on PTSD GWAS-derived gene associations. Each gene was weighted by its MAGMA Z-score and inversely weighted by its gene-specific technical noise level in snRNA-seq data from Tran et al.^66^ This process generated cell-specific raw disease scores. Additionally, we computed 1,000 sets of cell-specific raw control scores from matched control gene sets, ensuring similarity in gene set size, mean expression, and expression variance with the putative disease process genes. Subsequently, we normalized both the raw disease scores and raw control scores for each cell, yielding normalized disease scores and normalized control scores. The computation of these scores utilized the default settings of the scDRS compute-score function. Finally, we conducted cell-type-level assessments to link 16 cell-types to disease and explore heterogeneity in disease association across cells within each cell-type.

Finally, to evaluate the effects of the four PTSD risk-associated SNPs in the credible set of the FOXP2 locus on *FOXP2* expression, we analyzed bulk RNA seq data from medial amygdala of 182 individuals (106 neurotypical controls and 76 individuals diagnosed with PTSD) generated by Jaffe et al.^70^ Data processing was conducted using SPEAQeasy.^71^ Genes with low expression were filtered out with edgeR’s filterByExpr function,^72^ followed by library size normalization and voom transformation. Cell type proportions were estimated with Bisque^73^ using the data generated by Tran et al^66^ as reference.

We tested the effect of each of the four SNPs and inhibitory cell proportion as additive and interactive effects on *FOXP2* expression following a prior approach.^74^

Model fitting was performed adjusting for SNP dosage (coded as 0, 1, 2), the proportion of inhibitory neurons and their interaction, as well as diagnosis (PTSD vs. neurotypical control), sex, age at death, RNA integrity number (RIN), estimated proportions of astrocytes and excitatory neurons, oligodendrocytes, microglia, OPCs, and T cells, the proportion of reads mapped to the mitochondrial chromosome, and the first two ancestry principal components.

For visualization, we regressed out all covariates except SNP dosage, inhibitory neuron proportion, and their interaction term, and plotted the resulting residualized values of *FOXP2* expression.

### Animals

Adult male c57BL/6 mice were obtained from Jackson Laboratory (Bar Harbor, ME). Animals were housed in groups of 4 in a temperature and humidity-controlled vivarium with a 12 hour on/12 hour off light cycle. All experiments were performed during the light stage. Except for the time spent in experimental chambers or during surgery, animals had *ad libitum* access to food and water. Animals were randomized to intracranial injection conditions (Foxp2 shRNA virus vs scrambled control virus), such that half of animals in each cage received the experimental and the other half received the control viral construct. Sample size was determined based on effect sizes from previously published experiments and in our laboratory, and no *a priori* sample size calculation was performed for this study. All experimental procedures were approved by the McLean Hospital Institutional Animal Care and Use Committee and are following the National Institutes of Health (NIH) Guide for the Care and Use of Laboratory Animals.

### Intracranial viral injections

Mice were anesthetized with inhaled isoflurane in preparation for brain surgery/cannulation, with subcutaneous meloxicam (5mg/kg) for analgesia. Coordinates for targeting the bilateral medial intercalated cell cluster were as follows: medial-lateral +/-3.0, anterior-posterior −1.06, dorsal-ventral −5.2 as measured in mm from the interpolated bregma. Each mouse received bilateral injections of either Foxp2 shRNA or scrambled control. The following viral constructs were used, both in AAV9 vectors (Charles River, Rockville MD): ITR-U6-Foxp2shRNA-Ef1α-GFP-PolyA-ITR and ITR-U6-shRNA(Scramble)-Ef1α-GFP-PolyA-ITR. The Foxp2 shRNA sequence was 5’-GCAACAGTTCAATGAATCAAA-3’, as previously described.^75^ Following surgery, each mouse was allowed to recover in a separate cage on a heating pad to maintain body temperature. After consciousness was attained, the mice were group housed. The virus was allowed to express for 4 weeks with mice in home cages, prior to initiation of behavioral assessments.

### RNAScope in situ hybridization, Imaging, and Image Analysis

Adult (8-12 weeks) mice were anesthetized with isoflurane and humanely killed by rapid decapitation. Brains were dissected, immediately fresh-frozen on dry ice, and kept in −80C until further processing. Brains were sectioned into 16 µm slices through the entire region of the amygdala, placed on positively charged slides, and stored at −80C until further processing. Subsequent fixation in 4% paraformaldehyde and *in situ* hybridization was performed as described by the manufacturer (RNAscope Multiplex Fluorescent Reagent Kit v2, Advanced Cell Diagnostics, Newark CA) with one change: digestion time with Protease IV was 15 minutes instead of 30 mins, as described in the User Manual. RNA probes for *Foxp2, Oprk1,* and *Gal* were purchased from Advanced Cell Diagnostics.

Fluorescent Imaging was acquired on Leica TCS-SP8 scanning confocal microscope equipped with LAS-X software, v.3.5.7.23225 (Leica Microsystems Inc, Buffalo Grove IL). Laser power, number of z-slices, and image gain were kept consistent within each cohort. Slices in which the amygdala could not be fully visualized were excluded from the final analysis. Fluorescence area was calculated with ImageJ software (National Institutes of Health, Bethesda, MD). Statistical analysis and graphing were performed using GraphPad Prism (Boston, MA). Illustrations were created with BioRender.com.

### Behavioral Procedures

A battery of behavioral tests was performed ∼4-5 weeks after viral injections. Open Field (OF) test, Elevated Plus Maze (EPM), and Novel Object Recognition tests were performed on sequential days. Acoustic startle was performed before fear conditioning in one cohort, and after fear conditioning, in a separate cohort of male mice.

Open Field Test: The open field test was used to characterize locomotor activity in a novel environment. It was carried out in the open field arena (a cubic box with 50×50×50cm). The top of the box was uncovered, and the room was dimly illuminated (20 lux) to minimize anxiety effects on locomotion. All mice were placed into a corner of the box at the beginning of the trial and allowed to explore the box for 10 min. The traveled distance and zone time was tracked using Ethovision 14.5 software (Noldus, Wageningen, the Netherlands).

Elevated Plus Maze Test: The elevated plus maze test was used to assess anxiety-related behavior. The experiment was carried out in a chamber elevated 70cm above the ground that consisted of two open arms (30 × 5 cm), two closed arms (30 × 5 × 15 cm), and a central square (5 × 5 cm) with an open top. For testing, animals were placed at the central square of the maze facing the closed arm and were allowed to freely explore the maze for a duration of 10 min under constant low illumination conditions (20 lux). The traveled distance and zone time was tracked using Ethovision 14.5 software (Noldus, Wageningen, the Netherlands).

Novel object recognition: Novel object memory test was performed in a circular open field chamber with an open top (46 cm diameter, 38 cm height). For each mouse, this test was performed in four rounds, each consisting of a 5-minute exploration period, followed by a 5-minute intertrial interval. The mice were presented with a total of three white, polyactic acid objects: a cone, a cylinder with a rounded top, and a cylindrical star. During the first three rounds, the mouse was allowed to explore the cone and the cylinder with the rounded top, which were alternated between two locations in each round. During the fourth round, one of the objects was switched with a novel object (cylindrical star), and the mouse was allowed to explore as above. The chamber and objects were cleaned with dilute soap solution between trials. Object exploration was defined as a nose entering a 2 cm zone from the edge of each object. Zone time was tracked using Ethovision 14.5 software (Noldus, Wageningen, the Netherlands).

Acoustic Startle: Acoustic Startle was performed in 4 identical startle chambers (6×6×5 inches) with metal rod flooring attached to a load-cell platform in a sound-attenuating chamber (Med Associates, Georgia, VT). Mouse response to a burst of white noise resulted in a displacement of a transducer in the platform, with voltage amplification and digitization to arbitrary units by an analog-to digital converter card. Startle stimuli were generated by an audio stimulator (Med Associated, Georgia VT). In cohort 1, mice underwent acoustic startle prior to acoustic fear conditioning, and in a separate cohort 2, mice underwent the same acoustic startle protocol, but on the day following fear conditioning. In both cohorts, mice underwent a 5-minute adaptation period following a habituation trial to 20 presentations of 90dB startle stimuli, presented in 30 sec intervals. The mice were then presented with 20 stimulus trials, each trial at 7 different startle intensities (60dB, 70dB, 80dB, 90dB, 100dB, 110dB, 120dB), every 30 seconds. In each trial, the startle stimuli were presented at randomized order. Experimental and control mice were alternated between four acoustic startle chambers, such that no one group was consistently measured in one set of chambers. A dilute soap solution with a novel scent was used for cleaning. Startle amplitude is expressed in arbitrary units, with the mean averaged across the 20 trials for each of the seven startle-eliciting intensities. Startle data were analyzed using a two-way ANOVA with condition (*Foxp2* KD vs scrambled control) as a between-subjects factor and startle intensity (60, 70, 80, 90, 100, 110, 120 dB) as a within-subjects factor.

Fear Learning: Male mice, living in group-housed home cages, were habituated to the fear conditioning chambers for 15 minutes for two consecutive days prior to the start of behavioral testing. The chambers (30.5 x 25.9 x 30.5 cm, Lafayette Instruments) had a shock grid floor, were lit by white house lights, and were cleaned with Quatricide. On the day of fear learning, mice were allowed to explore the chamber for 180 sec prior to being exposed to five randomly spaced trials of tone-shock pairings of 30 s, 75 dB, 6 kHz tone-conditioned stimulus that co-terminated with a 1sec, 0.6 mA shock-unconditioned stimulus administered through the grid floor (Actimetrics, Wilmette IL). Experimental and control mice were alternated between four fear conditioning chambers, such that no condition was overrepresented in a single chamber. For subsequent RNA sequencing and qPCR analyses, animals were humanely killed by brief anesthesia with inhaled isoflurane followed by rapid decapitation 2 hours after fear learning. For subsequent Extinction and Extinction retention testing, mice were returned to their group-housed home cages for 24 hrs prior to the start of Extinction. Extinction and Extinction retention were performed in a different context (smooth floors, cages cleaned with 70% ethanol, red house lights in an otherwise dark room). Extinction training consisted of 30-tone CS presentations (30-s tone, 6 kHz, 75 dB, with no shocks) separated by a 60 s intertrial interval. Extinction retention (15 presentations of 30-second, 6kHz, 75dB tones, with 60s intertrial interval) was performed 24 hours after extinction training. Freezing analysis was performed with FreezeFrame5 software (Actimetrics, Wilmette IL) and was used to analyze freezing levels, with the freezing threshold chosen as follows: all thresholds started at 1.0 and were individually adjusted for each chamber until FreezeFrame5 correctly identified three distinct 10s intervals as either freezing or movement episodes. Freezing was defined as lack of any observed movement except for the muscles of respiration. The experimenter performing the thresholding was blind to the groups (Foxp2 shRNA vs scrambled shRNA injected mice) analyzed. Experimental groups were counterbalanced with controls during fear learning, acoustic startle, open field, elevated plus maze, novel object recognition, RNA extraction, cDNA synthesis, and qPCR.

### Intraperitoneal Lithium administration

Lithium Chloride (100mg/kg) or saline control was delivered intraperitoneally (I.P.) 30 mins prior to fear learning, as previously described.^76^ Following the auditory fear conditioning protocol described above, mice were humanely killed by brief anesthesia with inhaled isoflurane followed by rapid decapitation and brains were processed for RNAScope.

### RNA isolation and cDNA construction

Mice were humanely killed by brief isoflurane anesthesia followed by rapid decapitation at 5 weeks after viral-mediated knockdown of *Foxp2* (or scrambled control). Brains were collected, immediately fresh frozen, and stored at −80C until further processing. Tissue punches (1mm internal diameter, 1mm depth) were collected from the bilateral amygdala centered just above the mITCs to avoid the medial amygdala, where Foxp2 is also expressed. To reduce bias, punches were taken from alternating experimental vs control samples. Total RNA was isolated and purified using the Total RNA Purification Kit (Norgen Biotek Corp, Thorold, ON, Canada) according to the protocol provided by the manufacturer. RNA concentration was measured with Qubit 2.0 Fluorometer (Thermo Fisher Scientific) and reverse transcribed into cDNA with the Super Script IV First-Strand Synthesis System (Thermo Fisher Scientific) according to manufacturer instructions. cDNA was stored in −30C until further processing.

### qPCR

cDNA was amplified on an Applied Biosystems ViiA7 Real-Time PCR System with Power SYBR Green PCR Master Mix (Thermo Fisher Scientific). Gapdh was used as a control (Gapdh-fwd 5’ TATGACTCCACTCACGGCAA 3’, Gapdh-rev 5’ ACATACTCAGCACCGGCCT 3’). Subsequent data were analyzed using the ΔΔCt method.^77^ The following primers were used: Foxp2 (Foxp2-fwd 5’ GACCACATCGACAGCAATGG 3’, Foxp2-rev 5’ CTGCAATCACGGGTTCTTCC 3’), Pbx3 (Pbx3-fwd 5’ GCCAAATTGACCCAGATCAGA 3’, Pbx3-rev 5’ CGGAGAAGGTTCATCACATGT 3’), Tac2 (Tac2-fwd 5’ AGGGAGGGAGGCTCAGTAA 3’, Tac2-rev 5’ CCTTCCAGAGAGACAGGGC 3’), Slc12a3 (Slc12a3-fwd 5’ AAGTTCACGTCCTTCCCGAC 3’, Slc12a3-rev 5’ GGAAGGGTGCAATCATGTCC 3’), Mesp2 (Mesp2-fwd 5’ AGACTGGACACTGGACACAA 3’, Mesp2-rev 5’ CAGGGACAGGCAGGGTTC 3’), Amh (Amh-fwd 5’ TGGACACCATGCCTTTCCC 3’, Amh-rev 5’ AGGGTCTCTAGGAAGGGGTC 3’).

### RNA Sequencing

Bulk sequencing of cDNA from amygdala punches with polyA selection was performed by Genewiz Next Generation Sequencing (Azenta Life Science, South Plainfield, NJ). RNA integrity and library size were assessed with an Agilent Tapestation. Sequencing was performed on Illumina platform, 2×150bp, with a depth of 50 Million paired end reads per section. The mean quality score per sample was 38.30, with 91.88% of bases >=30.

### Transcriptional analysis

FASTQ files were processed with SPEAQeasy^71^ to extract features at the gene, exon, junction, and transcript levels. Using the SPEAQeasy-aligned data, genes with low expression were filtered using edgeR’s filterByExpr function^72^, followed by library size normalization, voom transformation, and model fitting with eBayes.^78^ Because we controlled for all relevant covariates (all mice were of male sex, behavioral testing was performed in one batch, and experimental/control samples were alternated during punching and RNA extraction) in the experimental design, the final model included only the *FOXP2* knockdown variable.

### Pathway Enrichment Analyses

Gene set enrichment analysis was performed using the fgsea package with Reactome pathway annotations.^79^ Genes were ranked by their log fold change and normalized enrichment scores (NES) were computed to identify significantly enriched pathways.

### Electrophysiology recordings in amygdala slice preparation

To examine how *Foxp2* knockdown alters the activity of amygdala intercalated neurons, we performed electrophysiology recordings in slice preparations from scrambled control vs *Foxp2* knockdown mice. Mice were anesthetised with isoflurane and transcardially perfused with an ice-cold N-methyl-D-glucamine (NMDG) solution of the following composition (in mM): 93 NMDG, 2.5 KCl, 1.2 NaH2PO4.H2O, 30 NaHCO3, 20 HEPES, 25 D-glucose, 5 sodium ascorbate, 3 sodium pyruvate, 2 thiourea, 10 MgSO4.7H2O, and 0.5 CaCl2.2H2O. After rapid decapitation, brains were extracted and cut into 250-µm coronal sections containing the amygdala using a vibratome (Leica VT1000S) in the same ice-cold NMDG solution. Slices were briefly placed in a warm NMDG solution (32°C) for 10-12 min, before being transferred to a long-term holding chamber containing artificial cerebral spinal fluid (ACSF) (in mM): 126 NaCl, 1.6 KCl, 1.1 NaH2PO4, 1.4 MgCl2, 2.4 CaCl2, 26 NaHCO3, and 11 glucose, osmolarity ∼290 mOsmol l^−1^, and pH 7.3 with KOH (32–34°C). Before recording, sections were transferred to the recording chamber containing aCSF maintained at 32°C. Throughout slice preparations, all NMDG and aCSF solutions were continuously saturated with 95% O2/5% CO2. ITC neurons were visualized on an upright microscope (model BX51WI- Olympus).

Whole-cell recordings were made using a MultiClamp 700B amplifier (Axon Instruments, Union City, CA) and collected with Clampex10.6. ITCs were located ventral to the internal medial and external (lateral) basolateral amygdala capsule. Furthermore, this cluster was identified by GFP fluorescence, as well as electrophysiological properties that have been previously characterized.^80^ Specifically, we find a small mean membrane capacitance (scrambled control Cm=38.84 ± 9.456), and large mean input resistance (scrambled control Rin= 374.3 ± 96.93 MW). For cellular activity experiments, recording electrodes (3-5 MΩ) were filled with (in mM) 130 potassium-D-gluconate, 10 KCl, 10 HEPES, 0.5 EGTA, 10 sodium creatine phosphate, 4 Mg-ATP and 0.3 Na2GTP. Immediately after breaking into the cell, cells were switched into current-clamp mode, and the resting membrane potential was recorded. Membrane potential for each neuron was set to −70 mV by current injection via the patch amplifier before switching to voltage-clamp to record action potentials. A current step protocol consisting of 10 steps (−25 to + 200pA, 25pA increments, 400ms in duration, three seconds apart) was applied and the number of action potentials at each step was recorded. Access resistance was monitored throughout recording, and cells were only accepted for analysis if the initial access resistance Ra was less than 20 MW and did not change by more than 20% throughout the recording.

### Human Postmortem Brain Differential Expression Analysis

We used data from the Jaffe et al^70^ published RNA-seq dataset to test the effect of PTSD risk SNPs on FOXP2 gene expression. To approximate the mouse experimental design, we created a binary variable representing low versus high *FOXP2* expression using the median as the threshold. Model fitting was performed using eBayes,^78^ adjusting for: binary *FOXP2* levels, diagnosis (PTSD vs. neurotypical control), sex, age at death, RNA integrity number (RIN), estimated proportions of astrocytes, inhibitory and excitatory neurons, oligodendrocytes, microglia, OPCs, and T cells (calculated with Bisque using the data generated by Tran et al.^66^ as reference), the proportion of reads mapped to the mitochondrial chromosome, and the first two ancestry principal components. We identified signals passing multiple testing correction using a False Discovery Rate (FDR)-adjusted p-value < 0.05.

### Transcriptional correlation of PTSD GWAS prioritized genes and genes significantly up/downregulated after Foxp2 KD

Starting with a list of prioritized PTSD putative causal genes from Nievergelt et al,^8^ we identified mouse orthologs of the prioritized genes that had an adjusted p-value < 0.01 after *Foxp2* KD and assigned the same gene prioritization category (Tier 1 or Tier 2) to the mouse orthologues. We excluded genes that had no defined mouse orthologues from the analysis.^81^ Tier 1 genes are defined as having a greater likelihood of being causal risk genes responsible for the disease signal, whereas Tier 2 genes are described as having a lower likelihood and the absence of a minimum level of evidence as a causal risk gene, albeit meeting a higher level of evidence than unranked genes.^82^ As a negative control for general CNS pathology in a neuropsychiatric illness, we also compared the *Foxp2* KD dataset to the prioritized genes in Alzheimer’s Disease GWAS that used the same prioritization methodology.^82^ After noting a marked decrease in mouse orthologues of PTSD Tier 1 genes and an increase in the expression of mouse orthologues of PTSD Tier 2 genes after *Foxp2* KD, we evaluated the relative contribution to the ranked tiering of individual components of the composite PTSD prioritization score.^8^ We mapped *Foxp2* KD up and downregulated mouse orthologues of the PTSD GWAS prioritized genes to the individual gene prioritization components from Nievergelt et al,^8^ including locus location on the human genome, highest posterior inclusion probability (PIP), SNPs with combined annotation depletion (CADD) scores >12.37, gene probability of Loss-of-function Intolerance (pLI), as well as Loss-of-Function Observed/Expected Upper Bound Fraction (LOEUF) score, an updated metric from gnomAD that measures a gene’s intolerance to mutation.^83^ Using the same list of mouse orthologues significantly up or downregulated after *Foxp2* KD, we mapped the available evidence for human orthologues of these PTSD risk genes across categories of evidence used for the prioritization, including positional mapping, eQTL mapping, and chromatin interaction (CI) mapping.

## Results

### Fine Mapping of the FOXP2 locus reveals variants associated with PTSD

To narrow down the GWAS-identified FOXP2 genomic locus and further identify potential causal variants driving the association with PTSD, we performed fine mapping of the *FOXP2* locus (*chr7:* 113858363- 114,290,415; **Figure 1A**), which contained 132 SNPs that passed GWAS-significance level (p<5×10^−8^), revealed four SNPs in a credible set (*chr7:* 114,056,055-114,071,035; rs12533005 - risk allele: G, rs2894699 - risk allele: C, rs7785701 - risk allele: G, and rs1476535 - risk allele: T).^8^ These PTSD risk-associated SNPs are highlighted in red in the right panel of Figure 1A.

To further identify subpopulations of cells in the human amygdala enriched for PTSD genomic risk pathways, we used MAGMA to perform gene set enrichment analysis. We identified ten cell types, including five inhibitory cell subpopulations, that were enriched for PTSD risk genes (**Figure 1B**) using sing, a human amygdala cell-type taxonomy derived from single-nucleus RNA sequencing (snRNA-seq) of neurotypical donors^66^. These included glial cell types, including microglia (p_adj_ =0.001), oligodendrocyte progenitor cells (p_adj_ = 0.001), astrocytes (p_adj_ = 0.007), and oligodendrocytes (p_adj_ = 0.02). Neuronal cells included inhibitory subtypes E (p_adj_ = 0.0005) (marked by expression of transcription factor *ONECUT2*), D (p_adj_ = 0.001) (marked by expression of *Kit* receptor tyrosine kinase and *TACR1* tachykinin receptor), F (padj = 0.001) (marked by expression of *LHX6* transcription factor), C (p_adj_ = 0.004) (marked by expression of *NPFFR2,* a neuropeptide receptor that is also associated with threat processing), and B (p_adj_ = 0.004) (marked by expression of *CALB2* calretinin and *ADRA1B* adrenergic receptor).

Using the same amygdala cell type taxonomy, we characterized FOXP2 transcriptional activity across cell types.^66^ Of these, inhibitory subtypes C and E, as well as microglia, were enriched for *FOXP2* expression (**Figure 1C**). When calculating PTSD risk relevance scores and accounting for single cell heterogeneity, subtype C was again one of the enriched inhibitory cell types (**Figure 1D**). Inhibitory cell subtype C is also characterized by expression of transcription factor *TSHZ1* which, like FOXP2, is both a marker of the ITCs and a critical mediator for the migration, differentiation, and survival of ITCs during development, suggesting that this cell type is, at least partially, composed of the ITC population.^84^

Due to their scarce location between amygdala subregions, dissection of ITCs from postmortem human brains is not technically feasible. In the absence of ITC tissue from postmortem brains, we next examined the effects of the four causal PTSD risk SNPs at the *FOXP2* locus on *FOXP2* expression in the human postmortem medial amygdala (MeA) – a *FOXP2-*expressing region that neighbors the ITCs. We observed significant associations for all SNPs in the credible set. Specifically, carriers of the PTSD risk alleles for *FOXP2* exhibited decreased *FOXP2* expression (ie, all estimates were negative; rs12533005: β = - 0.57, p = 0.03781; rs2894699: β = - 0.57, p = 0.0389; rs7785701: β = - 0.57, p = 0.03781; rs1476535: β = - 0.57, p = 0.04023).

Conversely, in the analyses including rs7785701 and rs1476535, samples with relatively higher proportions of inhibitory neurons were associated with increased *FOXP2* expression (p = 0.00824 and p = 0.00838, respectively; **Figure 1E**). This overall positive relationship was driven by the homozygotes of the risk alleles, while the heterozygotes or the homozygotes of the non-risk alleles significantly differed, showing the opposite relationship (rs7785701: GG: r = 0.263 vs. CG/CC: r = −0.107 - Fisher r-to-z = 2.45, p = 0.0143; rs1476535: TT: r = 0.274 vs. CT/CC: r = 0.111 - Fisher r-to-z = 2.53, p = 0.0114). These data suggest that the top potentially causal *FOXP2* SNPs are associated with differential *FOXP2* mRNA expression in the human amygdala and suggest dysregulated signaling in ITC cell populations that are associated with amygdala regulation of threat processing.

### Transcriptional regulation of Foxp2 after fear learning in the mouse amygdala

Given substantial enrichment of *FOXP2* in inhibitory cells characterized by expression of *TSHZ1* ITC marker, we next characterized the role of *Foxp2* in the ITCs using a mouse model of fear/threat learning. Dynamic regulation of genes with a vital role in learning and memory has been observed in numerous amygdala-expressed pathways, including mTOR, Wnt, and BDNF pathways, among others.^41,85,86^ To test the hypothesis that *Foxp2* mRNA in the ITCs is similarly dynamically regulated, we examined the levels of *Foxp2* mRNA immediately after or 8 hours after fear conditioning (**Figure 1F**). Compared to baseline levels in control mice, we find that *Foxp2* is decreased immediately after fear learning of tone/shock pairings, with mRNA levels returning to baseline levels at 8 hours after fear conditioning **(Figure 1G, I).** These data suggest dynamically fluctuating *Foxp2* levels may represent a functionally necessary molecular process for the formation or expression of fear memories in the amygdala.

### Fear expression is reduced after Foxp2 KD in the amygdala ITCs

ITCs have been previously defined as critical cellular mediators of fear extinction in the amygdala;^87^ However, the role of the Foxp2 transcription factor in fear memory remains unknown. To test the role of Foxp2 in acquisition and expression of fear learning, we sought to decrease expression of *Foxp2* in the medial ITC cell cluster, bilaterally, using an shRNA expression AAV virus, followed by behavioral characterization after fear learning.

Notably, several nomenclatures of ITC clusters in rodents have been proposed. Here, we use the most recent nomenclature based on a study of detailed anatomical connectivity of the ITCs in mice, with a focus on the role of *Foxp2* in the main ITC cluster (mITC) (**Figure 1H**), which projects to the central, medial and cortical amygdala, with considerable projections to the other ITCs.^64^ To knockdown *Foxp2*, we injected AAVs containing short hairpin RNA (shRNA) targeting *Foxp2* under control of a U6 promoter, followed by auditory fear conditioning (5 tone-shock pairs in context A), fear expression / extinction training (30 tones without shocks in context B), and extinction retention (15 tones without shocks in context B, with each behavioral session separated by 1 day). (**Figure 2A)** This viral shRNA construct has been previously shown to decrease Foxp2 protein levels in the mouse striatum.^75^ Based on previous studies of the role of ITCs in extinction, we expected to observe a deficit in fear extinction 24 hours after fear acquisition.^88^ Contrary to our hypothesis, mice with *Foxp2* KD displayed a robust and significant decrease in fear expression during both fear acquisition and fear extinction training (**Figure 2B**). A two-way ANOVA for fear acquisition revealed a significant effect of condition group (F_(1,54)_=36.85, p<0.0001) and trial (F_(5,270)_=52.26, p<0.0001). *Post hoc* Tukey’s multiple comparison test revealed no difference in the 30 second interval immediately preceding the first tone (pre-CS, p=0.98) or during the first 30 second tone which terminated with a shock in the 29-30 sec interval (Trial 1, p=0.89). The subsequent four trials showed a significant decrease in freezing in the *Foxp2* KD condition (Trial 2, p=0.0014; Trial 3, p<0.0001; Trial 4, p<0.0001; Trial 5, p<0.0001). A two-way ANOVA for extinction training (i.e., fear testing) revealed a significant effect of condition group (F_(1, 245)_ = 41.87, p<0.0001) and trial ((F_(6, 245)_ = 7.113, p<0.0001), but no group by trial interaction (F_(6, 245)_ = 1.609, p=0.1451). Šídák’s multiple comparisons test revealed no difference in the 30 second interval immediately preceding the first tone (pre-CS, p=0.99) or during the first through fifth tones (Trial Bin 1, p=0.1146). The subsequent five trial bins showed a significant decrease in freezing in the *Foxp2* KD condition (Trial bin 2, p= 0.0371; Trial bin 3, p< 0.0125; Trial bin 4, p< 0.0415; Trial bin 5, p= 0.0081, Trial bin 6, p= 0.0175). No significant differences between the experimental and control groups were observed on the Extinction Retention day (Two-way ANOVA revealed a significant effect of trial bin (F_(1.694, 35.58)_ = 4.193, p=0.0287), with no effect of experimental condition (F_(1, 21)_ = 1.232, p=0.2795) and no group by trial interaction (F_(1.694, 35.58)_ = 2.294, p=0.1229). Šídák’s multiple comparisons test revealed no difference in the 30 second interval immediately preceding the first tone (pre-CS, p=0.9998) or during the trial bins (Trial bin 1, p= 0.9998; Trial bin 2, p= 0.2249; Trial bin 3, p=0.9018) (**Figure 2B**). These data suggest that *Foxp2* KD results in robust decrease in fear acquisition and expression during extinction training, and that by the time of extinction retention the control animals had already achieved a floor level of low freezing similar to the experimental group.

**Figure 2.**
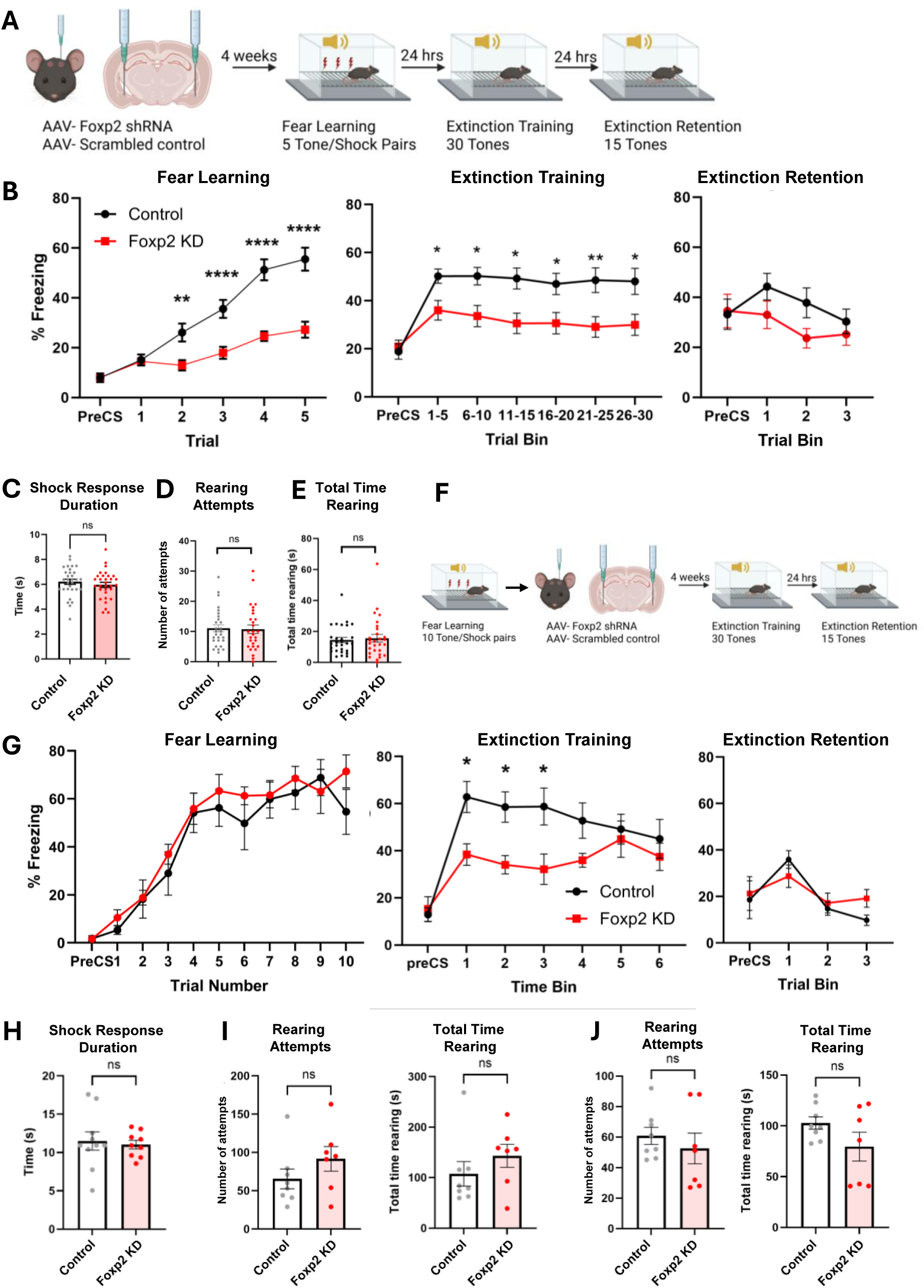
Expression of fear learning is decreased after *Foxp2* KD in the amygdala. (A) Foxp2 shRNA or scrambled control constructs were injected bilaterally targeting the mITC. Following viral expression, male mice underwent fear conditioning with 5 tone/shock pairs (context A), followed by 30 tone extinction (context B) and 15 tone extinction retention (context B). B. Foxp2 KD (n=28) significantly reduced fear expression during fear learning and extinction training as compared to mice injected with scrambled shRNA control (n=28), without an observed difference in (C) duration of behavioral response to the shock, (D) number of rearing attempts, or (E) total time rearing during fear learning. (F) To test the effect of Foxp2 KD on extinction training and expression, male mice underwent fear conditioning with 10 tone/shock pairs prior to Foxp2 KD (red, n=7) (or scrambled control, black, n=8) targeting bilateral mITCs, followed by extinction training and extinction retention, as above. (F) Allowing 4 weeks for viral expression, mice demonstrated impaired expression of fear during extinction training, without an effect on extinction recall (G), duration of responding to shock stimuli (H), and without a change in the number of rearing attempts or total time rearing during extinction training (I) and extinction retention (J). 2-way Anova followed by multiple comparison testing. ****p<0.0001, ***p<0.001, **p<0.01, *p<0.05; two-way Anova followed by Tukey’s multiple comparison test (panel B) and ns, not significant, unpaired t-test (panels C-E, H-J).

To test the possibility that decreased freezing response was due to diminished processing of momentary pain from the footshock, we quantified the amount of time spent during a characteristic immediate shock response. Following a footshock, mice exhibit a stereotyped flight response of jumping followed by darting about the cage.^89^ We do not observe this response in the presence of an auditory cue alone, making this flight-like behavior a marker of physical response to a momentary footshock. To quantify this response, we measured the total amount of time spent during jumping and darting behavior in response to the unconditioned footshock and found no difference between the *Foxp2* KD and scrambled control groups (Unpaired t-test, p=0.3962), suggesting that pain sensitivity was not affected by the KD (**Figure 2C**). Next, we tested whether the observed decrease in freezing was due to decreased anxiety-like behavior by measuring rearing activity during fear learning. Rearing is a stress-sensitive behavior in rodents and has been used as a marker of increased anxiety-like behavior during stress.^90^ We observed no change in number of rearing attempts (Unpaired t-test, p=0.8612), or total time rearing (Unpaired t-test, p=0.7154) between the *Foxp2* KD and control conditions, suggesting the decrease in freezing behavior was not due to change in anxiety-like behavior or general locomotion levels during fear learning or expression (**Figure 2D, E**).

### Extinction expression is decreased after Foxp2 KD

The observed decrease in freezing during fear expression and extinction training could be due to: 1) a deficit in physical expression of fear learning, 2) an initial fear learning or fear consolidation deficit, in which an acoustic tone is not associated with a shock, or 3) decreased fear expression following fear consolidation. To differentiate between these possibilities, we performed fear conditioning prior to viral injection with Foxp2 shRNA, ensuring that all experimental and control animals had acquired and consolidated the initial fear learning and were physically capable of physical expression of fear (freezing) prior to *Foxp2* KD (**Figure 2F**). Mice underwent fear conditioning with 10 tone/shock pairs and demonstrated no difference in freezing levels among animals subsequently randomized to the scrambled control vs *Foxp2* shRNA groups (**Figure 2G**). Approximately 4 weeks after viral injection, fear extinction training (i.e., delayed fear expression) was performed through a presentation of 30 tones. Again, we found that *Foxp2* KD mice demonstrated significantly less freezing than the scrambled control-injected counterparts. A two-way ANOVA revealed a significant effect of group (F_(1, 91)_ = 18.95, p<0.0001) and time (F_(6, 91)_ = 7.780, p<0.0001), but no group by trial interaction (F_(6, 91)_=1.682, p=0.1343). Šídák’s multiple comparisons test revealed no baseline change during the 30 seconds prior to the first tone (PreCS, p=>0.9999); however, the first three time bins had significantly increased freezing in the control vs the *Foxp2* KD conditions (Time bin 1, p=0.0478, Time bin 2, p=0.0462, Time bin 3, p=0.0235; **Figure 2G**). We also tested extinction recall 24 hours after initial extinction training and found a significant effect of time (F_(15, 208)_ = 5.321, p<0.0001), but no group (F_(1, 208)_ = 0.6064, p=0.4370) or group by trial interaction (F _(15, 208)_ = 1.599, p=0.0762) (**Figure 2G**). We found no difference in the time responding to shock during initial fear conditioning (Unpaired t-test, p=0.7288) (**Figure 2H**), or the number of rearing attempts and total time rearing during extinction training (**Figure 2I**) and extinction retention (**Figure 2J**; Unpaired t-test, extinction training rearing attempts p=0.2286, total time rearing, p=0.3045; extinction retention rearing attempts p=0.4928, total time rearing, p=0.1691). Taken together, *Foxp2* KD mice showed decreased freezing during both initial fear learning and extinction training (delayed fear expression), suggesting a deficit in fear expression during repeated CS presentation. This pattern indicates that *Foxp2* KD affects fear expression, rather than initial fear acquisition or consolidation.

### Foxp2 KD in the mITC has no effect on exploratory behavior, novel object recognition, or acoustic startle

Although we observed a marked decrease in fear expression, we could not rule out that this phenotype is mediated by circuits specific to fear learning vs a generalized decrease in anxiety-like behavior. Lack of change in rearing activity during fear learning (**Figure 2D,E**) suggested no change in anxiety-like behavior; however, it did not rule out the possibility of decreased expression of such behaviors in the absence of acute stress. To test this possibility, we performed open field (OF) and elevated plus maze (EPM) tests in a subset of mice before fear learning (**Figure 3A**). We did not observe a change in percent of time spent in the center or periphery of the OF chamber (**Figure 3B**), time ambulating (**Figure 3C**), or distance travelled (**Figure 3D**). A two-way ANOVA for group (*Foxp2* KD vs scrambled control) by zone revealed a significant main effect of zone (F_(1, 120)_ = 1908, p<0.0001) and group by zone interaction (F _(1, 120)_ = 9.482, p=0.0026), without a significant main effect of group (F_(1, 120)_ = 2.928e-021, p>0.9999). Similarly, we observed no change in the percentage of time spent in the center, open, or closed arms of the EPM (**Figure 3E**). A two-way ANOVA for group by zone revealed a significant main effect of zone (F_(2, 117)_ = 3431, p<0.0001), but no significant effect of group (F_(1, 117)_ = 0.002628, p=0.9592), or group by zone interaction (F_(2, 117)_ = 2.29, p=0.1050).

**Figure 3.**
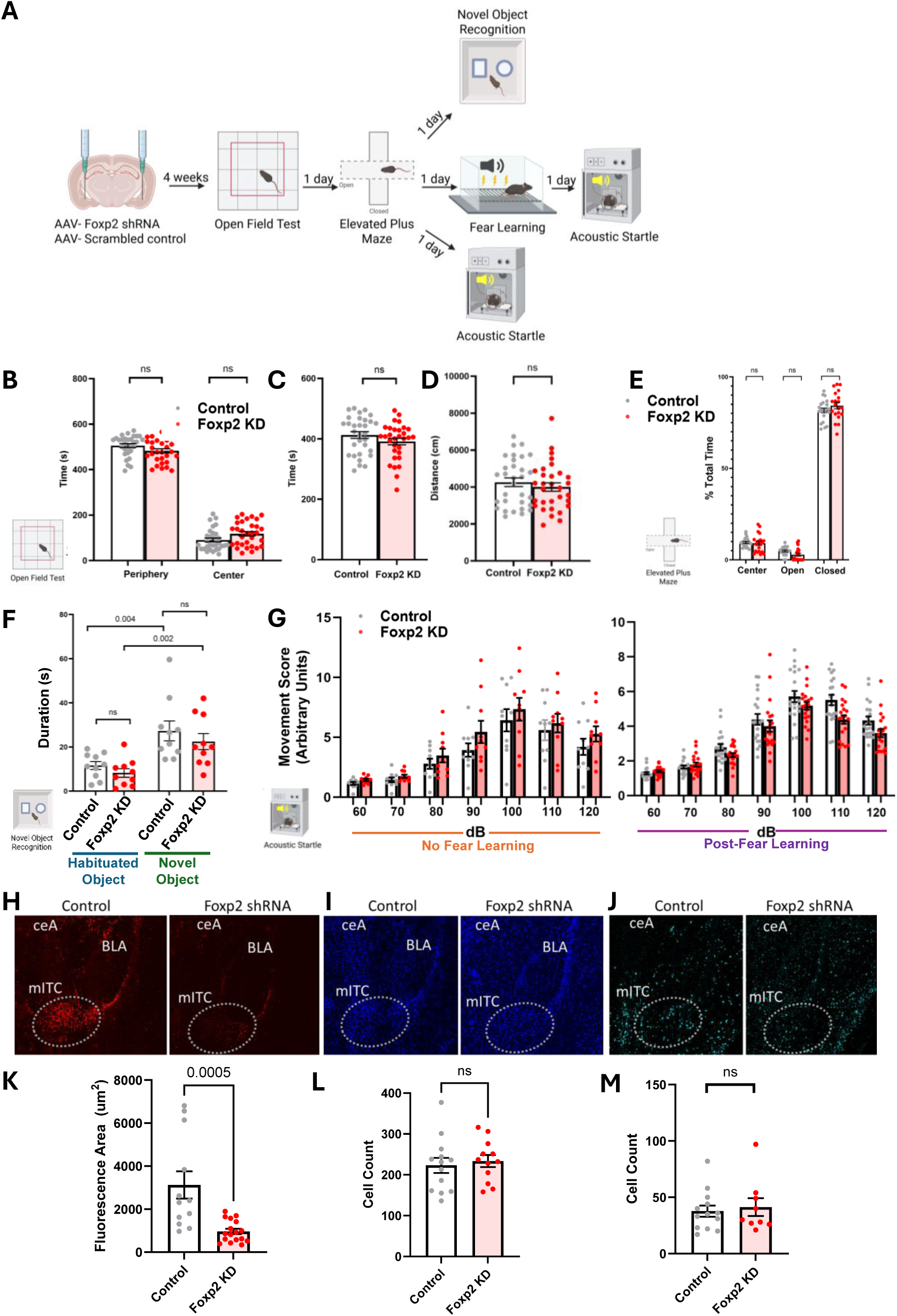
Transcriptional decrease in *Foxp2* has no effect on exploratory behavior, novel object recognition, acoustic startle, or ITC cell number and IEG expression. (A) Schematic of behavioral tests for exploratory behavior, declarative memory, and startle, in the setting of *Foxp2* KD. Adult male mice were separated into three cohorts after Elevated Plus Maze, with one cohort undergoing test for novel object recognition, one cohort undergoing fear learning followed by acoustic startle, and one cohort undergoing acoustic startle without fear learning. Open Field test shown as time spent in the center vs periphery (B), total time ambulating (C) and total distance travelled (D) shows no difference between *Foxp2* KD (red, n=29) and scrambled control (grey, n=31) injected mice. (E) Elevated plus maze shows no time difference spent in the center, open, and closed arms of the maze between Foxp2 KD (n=20) and control (n=21) injected mice. Two-way ANOVA followed by Sidak’s multiple comparison test. (F) Novel object recognition task demonstrates increased time exploring the novel object in both control (n=10) and Foxp2 KD (n=10) conditions. (G) Acoustic startle, reported as movement score in arbitrary units, showed no difference between the control and experimental conditions both in the absence of fear learning (Foxp2 KD, n=10; Scrambled control, n=10), and after fear conditioning (Foxp2 KD, n=10; Scrambled control, n=10), in a separate cohort of mice. Two-way ANOVA followed by Tukey’s multiple comparison test. RNAscope fluorescent *in situ* hybridization of *Foxp2* mRNA showing decreased fluorescence intensity of (H) *Foxp2* signal after Foxp2 shRNA-mediated knockdown, quantified as fluorescence area in (K); number of cells positive for DAPI labeling of the cell nucleus (I) quantified in (L), and (J) *Fos* mRNA, quantified in (M). Panels H-J are from the same sections of their respective experimental vs control conditions. Unpaired t-test, ns, not significant; n=14 amygdala (Control), 12 amygdala (*Foxp2* KD).

We next tested the possibility that a decrease in freezing during fear learning or expression represents a generalized decrease in learning or memory formation. In a subset of animals, we performed a novel object recognition task prior to fear learning (**Figure 3A**). Novel object recognition is a widely used spatial test for short and intermediate term memory.^91^ Following three trials of habituation to two objects in alternating positions, the mouse is introduced to a third, novel, object in place of one of the objects used during the training phase. The mouse is expected to preferentially explore the novel object; and the total time exploring the perimeter of the novel vs. habituated object is measured. We found a significant increase in time exploring the novel object in both the control and *Foxp2* KD conditions (**Figure 3F**). A two-way ANOVA for group by object zone (habituated vs novel object) revealed a significant effect of zone (F_(1, 36)_ = 22.47, p<0.0001), but no effect of group (F_(1, 36)_ = 1.671, p=0.2043) or group by zone interaction (F_(1, 36)_ = 0.05342, p=0.8185). Similarly to the finding in OF, we observed no difference in time ambulating during the novel object test (unpaired two tailed t-test, t_(18)_=1.642, p= 0.1180).

The Foxp2 transcription factor has a well-characterized role in vocal learning in a range of organisms, including songbirds, mice, and humans; ^92–94^ thus, an effect on auditory processing cannot be ruled out as a cause of our observed deficits in auditory fear expression. Our viral targeting of the mITCs is not expected to have passed through brain regions important for auditory processing, such as the thalamus or the primary sensory region of the cortex. Nevertheless, given FOXP2’s role in language learning, we next tested the possibility that the observed decrease in fear expression following *Foxp2* KD is due to a deficit in auditory processing such that mice may not hear the tone CS cue. To test auditory processing following *Foxp2* KD in the mITC, we performed an acoustic startle test in two separate cohorts of mice before and after auditory fear conditioning (**Figure 3A**). Following a period of acclimatization to the testing cage and habituation to a series of 90dB tones, mice were subjected to 20 trials of 7 bursts of white noise (60-120dB), presented in random order, with startle response measured in Arbitrary Units. In the pre-FC cohort, a two-way ANOVA of startle amplitude at each intensity (60, 70, 80, 90, 100, 110, 120 db) across the *Foxp2* KD and scrambled control group, showed a main effect of stimulus intensity (F_(6, 126)_ = 21.82, p<0.0001) and group (F_(1, 126)_ = 5.091, p=0.0258). However, Šídák’s multiple comparisons test revealed no group differences at any of the 7 stimulus intensities tested (**Figure 3G**). In the post-FC cohort, a two-way ANOVA showed a main effect of stimulus intensity only (F_(6, 126)_ = 22.46, p<0.0001), with no significant change in group (F_(1, 126)_ = 2.755, p=0.0994) or group differences at individual stimulus intensities (**Figure 3G**). This increase in startle response with higher dB white noise is the result expected in animals with intact auditory processing and is observed in both *Foxp2* KD and control groups, ruling out a general auditory deficit as a cause of fear expression deficits in *Foxp2* KD mice.

### Foxp2 KD has no effect on cell number or Fos immediate early gene activity in the ITCs after fear learning

To confirm the intended effect of our shRNA manipulation, we performed fluorescence *in situ* hybridization to examine *Foxp2* mRNA expression in the mITC in Foxp2 shRNA vs control injected mice. AAV delivery of Foxp2 shRNA resulted in a significant knockdown of *Foxp2* transcription (**Figure 3H and 3K**, unpaired t-test, p=0.0005). Given the magnitude of the reduction in fear learning after *Foxp2* KD and the essential role of the amygdala in auditory fear conditioning, we next tested whether AAV-induced decrease in cell viability in the amygdala contributed to the observed reduction in fear expression. Such deficits in fear learning had been previously described in amygdala lesion studies in rodents, and would be consistent with decreased freezing in response to auditory fear conditioning.^95^ We found no difference in DAPI-labeled cells in the ITCs after injection with Foxp2 shRNA vs scrambled shRNA constructs, suggesting that the observed fear expression deficit was not caused by generalized decrease in cell number or viability in the amygdala (**Figure 3I and 3L**, unpaired t-test, p=0.6657).

Another possibility for the observed decrease in fear expression is a general reduction in the neuronal response to fear conditioning. The amygdala is composed of ∼130 different cell types, a subset of which show a transcriptional response to fear learning by differential regulation of immediate early genes (IEGs) in response to stress.^96^ Within the intercalated cells, a subset of neuronal types has a robust IEG response to fear conditioning.^96^ We thus tested whether *Foxp2* KD results in a decreased transcriptional IEG response to stress, by labeling mRNA for IEG *Fos* in the ITCs two hours after fear learning. We found no difference in number of cells with *Fos* transcription between the Foxp2 shRNA and scrambled control conditions (**Figure 3J and 3M**, unpaired t-test, p=0.6915), suggesting IEG transcriptional response to stress is maintained after *Foxp2* KD, and the decrease in fear expression is likely mediated by yet unknown transcriptional mechanisms.

### Foxp2 KD increases intrinsic excitability of ITC neurons

To examine how *Foxp2* KD alters activity of amygdala intercalated neurons, we performed slice electrophysiology recordings from ITCs of *Foxp2* KD and scrambled control-injected mice using a GFP tag in AAV-expressing cells to identify neurons expressing *Foxp2* or scrambled control shRNA (**Figure 4A**). ITC neurons injected with *Foxp2* shRNA fired significantly more action potentials compared to scrambled-shRNA injected controls, with a significant main effect of input current on spike number (Two-way repeated measures ANOVA, main effect of current, F _(9, 180)_ = 126.2, p<0.0001), a main effect of viral condition (Two-way repeated measures ANOVA, F _(1, 20)_ = 12.95, p=0.0018), and a condition by current interaction (Two-way repeated measures ANOVA condition x current injected interaction F _(9, 180)_ = 6.636, p<0.0001) (**Figure 4B, D**). This finding was accompanied by an increased input-output slope (**Figure 4C**, unpaired t-test, p=0.0049). Consistent with increased ITC excitability, *Foxp2* KD neurons exhibited a lower firing threshold (**Figure 4E**, unpaired t-test, p=0.0086), decreased rheobase (**Figure 4F**, unpaired t-test, p=0.0011), and an increased action potential (AP) half-width (**Figure 4G**, unpaired t-test, p=0.0087). By contrast, there was no change in the action potential height (**Figure 4H**, unpaired t-test, p=0.6142), after-hyperpolarization height (AHP, **Figure 4I**, unpaired t-test, p=0.2347), or slope of depolarization (**Figure 4J**, unpaired t-test, p=0.8476), although the slope of repolarization was significantly reduced (**Figure 4K**, unpaired t-test, p=0.02).

**Figure 4.**
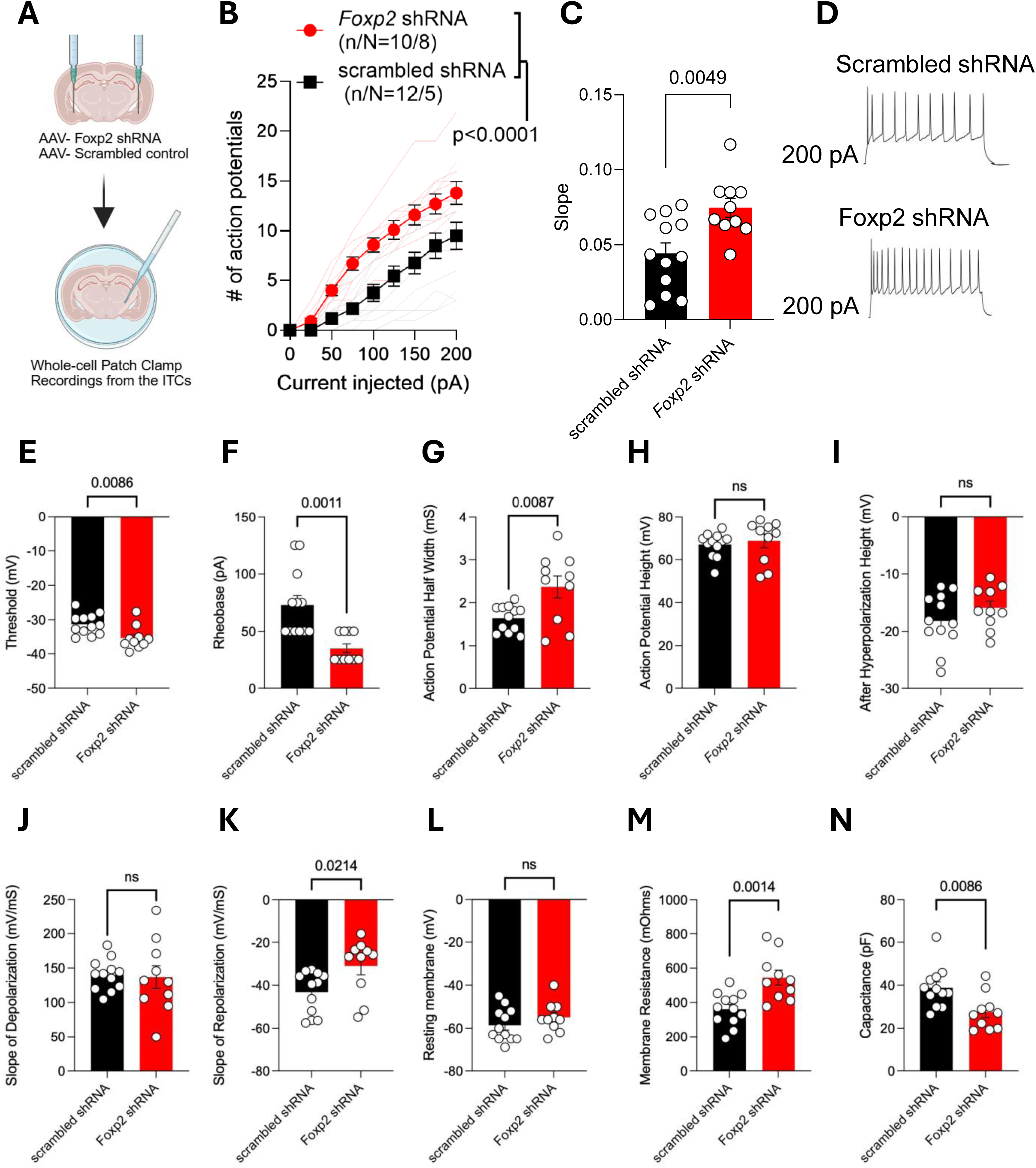
*Foxp2* KD increases intrinsic excitability of ITC cells. (A). Bilateral AAV injection (Foxp2 shRNA or scrambled control) targeting the mITC was followed by slice electrophysiology from ITCs expressing a viral tag. *Foxp2* KD (red) increases the number of (B) action potentials and (C) slope, p=0.0049 in the intercalated cells. (D) Representative traces for recordings of mice injected with scrambled shRNA or Foxp2 shRNA. (E) Threshold, p=0.0086 (F) Rheobase, p=0.0011 (G) Action potential half-width, p=0.0087, (H) Action potential height, p=0.6142, (I) After-hyperpolarization height, p=0.2347, (J) Slope of depolarization, p=0.8476, and (K) Slope of repolarization, p=0.0214. (L) Resting membrane potential, p=0.2597, (M) Membrane resistance, p=0.0014, (N) Capacitance, p=0.0086, n/N = number of cells/Number of mice. P-values in (C-N) are from unpaired t-tests.

At the cell membrane level, the resting membrane potential remained unchanged (**Figure 4L**, unpaired t-test, p=0.2597). However, *Foxp2* KD neurons exhibited an increased membrane resistance (**Figure 4M**, unpaired t-test, p=0.0014) and decreased capacitance (**Figure 4N**, unpaired t-test, p=0.0086).

Taken together, these findings indicate that *Foxp2* KD enhances ITC excitability through multiple complementary mechanisms. Neurons require less input to initiate firing (lower threshold and rheobase), respond more strongly to synaptic input (increased membrane resistance and reduced capacitance), and exhibit slower repolarization dynamics (reduced repolarization slope and increased action potential half-width). The reduced slope of repolarization and increased AP half width after *Foxp2* KD suggests that a decrease in K^+^ conductance could also, in part, mediate increased membrane resistance, as fewer open K^+^ channels would reduce leak conductance. Notably, AHP height, largely driven by K^+^ currents, was not altered, suggesting that the effects of *Foxp2* KD on excitability are unlikely to reflect a uniform suppression of all K^+^ dynamics, but rather a combination of changes across channel subtype.

Multiple potassium channels, including Kv4.2, expressed as the α-subunit of the A-type voltage-dependent potassium channel, and Kir3, encoding a family of protein-coupled inwardly rectifying potassium channels, are relatively enriched in the intercalated cells.^97,98^ The reduction in K^+^ channel efficacy could thus be mediated by a reduction in potassium channel transcription downstream of Foxp2 transcription factor signaling, which we examine in the following section.

### Transcriptional targets downstream of Foxp2 are involved in fear memory processes, synaptic signaling, and potassium conductance

Given the prominent role of Foxp2 in transcriptional regulation in the ITCs,^55^ we next examined the downstream transcriptional targets of Foxp2 in the mouse amygdala. FOXP2 regulon activity (genes from single cell sequencing data that are co-expressed with FOXP2/*Foxp2* and are putative direct binding targets based on *cis*-regulatory motif enrichment) is enriched in intercalated cells and contains the greatest number of genes of any other regulon set in the ITCs.^55^ As transcription factor regulation is activity dependent, we collected brains 2 hours after undergoing fear conditioning, to focus on downstream targets that are more likely to be involved in fear memory expression and consolidation. We performed bulk RNA sequencing on 1mm amygdala punches in mice injected with either *Foxp2* shRNA (n = 10 mice) or scrambled control (n = 9 mice; **Figure 5A**). Of the 19,510 differentially expressed genes (DEGs) in our dataset, 8,356 reached adjusted p-value<0.05, and 5,869 reached adjusted p-value <0.01 (**Figure 5C, Supplemental Data 1A**). Gene Set Enrichment Analysis (GSEA) after *Foxp2* KD identified an enrichment of downregulated genes in pathways involved in synaptic signaling, transcription of voltage gated potassium channels, and ligand binding, whereas upregulated genes were enriched in immune-related and metabolic pathways (**Figure 5E, Supplemental Data 1B**). Synapse-specific gene ontology analysis (SynGo)^99^ revealed an enrichment in regulation of trans-synaptic signaling, regulation of pre- and post-synaptic membrane potential, and synaptic vesicle cycle among the downregulated genes, and ribosomal translation enrichment among the upregulated genes (**Figure 5G, Supplemental Data 1C, D**).

**Figure 5.**
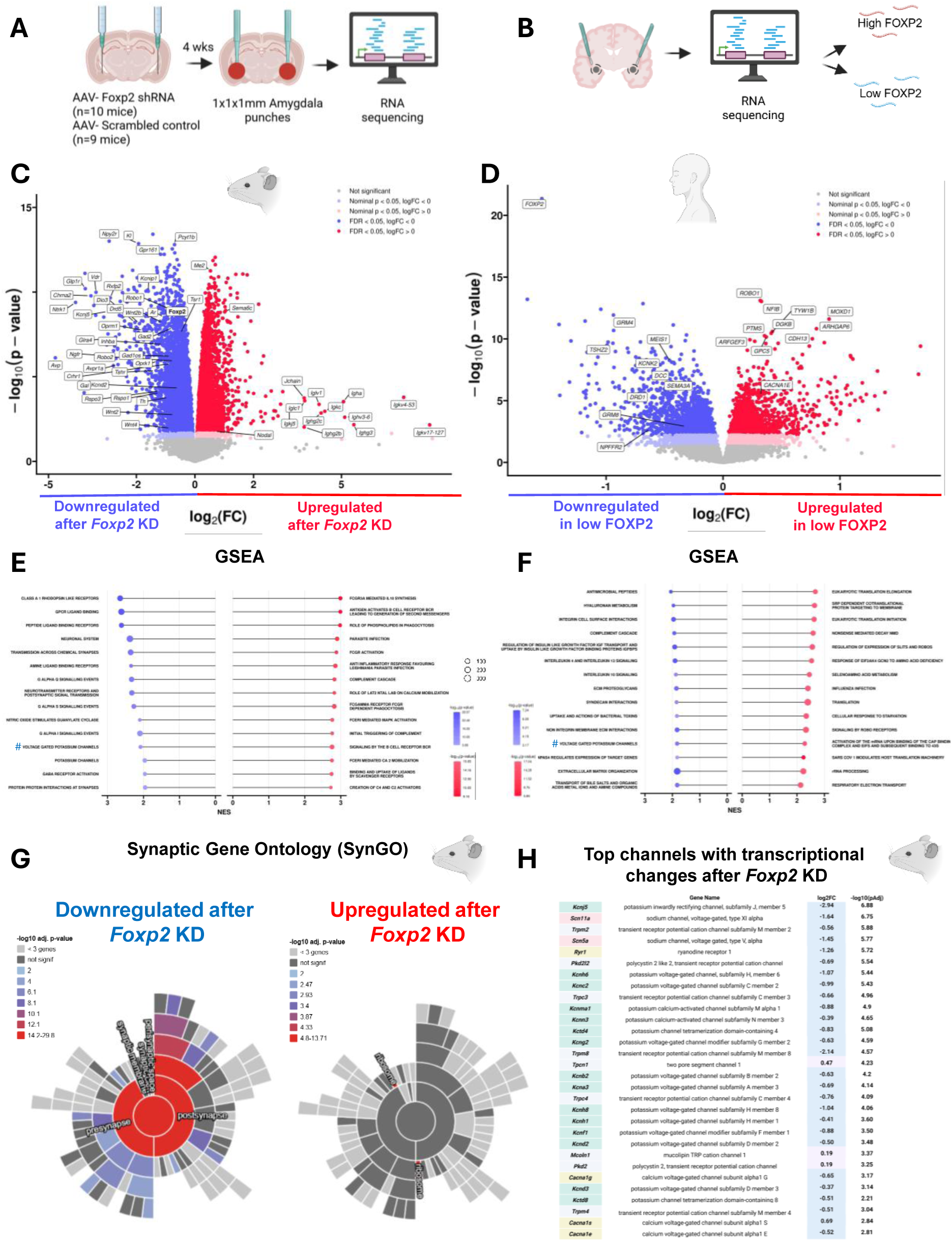
Foxp2/FOXP2 signaling regulates synaptic and fear-memory processes. Bulk RNA sequencing transcriptomic profiles following (A) bilateral *Foxp2* KD in the mouse mITCs and (B) genes associated with low-expressed *FOXP2* in postmortem human medial amygdala (MeA). Volcano plots of differentially expressed genes (C) after *Foxp2* KD in mice, and (D) genes showing significant transcriptional association with low *FOXP2* levels in the human MeA. Colored dots represent nominally significant genes (P < 0.05), with darker dots indicating those passing the p-adjusted threshold (P < 0.05). Gene set enrichment analysis (GSEA) of REACTOME pathways for *Foxp2* KD associated genes in mice (E) and low *FOXP2* associated genes in human postmortem dataset (F). The top 15 significantly upregulated and downregulated pathways (p-adjusted) identified by fgsea are shown. Normalized enrichment score, NES. Downregulated pathways are shown in blue, upregulated pathways are shown in red. Dot size is reflective of the number of genes in the respective dataset that is contributing to each pathway, while the color gradient indicates the log_10_(p-value). The pound sign (#) represents one of the top common downregulated pathways, voltage gated potassium channels. Synaptic gene ontology (SynGO) of p adjusted-significant genes after *Foxp2* KD in the mouse ITCs shows an enrichment among the downregulated, but not upregulated genes. Cellular component synaptic gene ontology is shown (G). Genes encoding potassium (mint green), sodium (pink), calcium (yellow), and mixed cation (grey) channels ranked by p-adjusted significance, are shown in (H).

Consistent with the electrophysiological finding of hyperexcitability after *Foxp2* KD and likely reduction in potassium channel activity, we found 64 significant DEGs for potassium, sodium, calcium, and mixed cation channels in our dataset^100^ (**Figure 5H, Supplemental Data 1E**). Of the 12 upregulated genes, six encode cation or calcium channels (*Pkd2, Mcoln1, Cacna1d, Trpv4, Tpcn2, Cacna1s*), four genes encode K+ channels (*Kcnj11, Kcnh2, Kcnj14, Kcnj8*), one gene encodes a mixed sodium/potassium channel (*Hcn2*), and one gene encodes a sodium channel (*Tpcn1*). The other 52 genes were downregulated and encoded 31 potassium, 4 sodium, 2 mixed potassium/sodium, and 15 calcium and cation channels (**Figure 5H**). Interestingly, two of the top downregulated potassium channels encode potassium channels that are relatively enriched in the ITCs - Kv4.2 (encoded by *Kcnd2*) and a channel of the Kir3 family (encoded by *Kcnj5)* (**Supplemental Data 1E**). Downregulation of these channels could, in part, explain the physiological hyperexcitability of the ITC neurons described above.

### Transcriptional correlation of genes with low FOXP2 expression in human postmortem medial amygdala

To further explore the hypothesis that transcriptional regulation of channels downstream of Foxp2 is mediating action potential propagation and cell excitability during fear learning, we examined transcriptional correlation of genes expressed in human postmortem medial amygdala (MeA) sections with high vs low *FOXP2* expression (**Figure 5B**). Of the 22,022 genes detected in this dataset, 5,005 reached an adjusted p-value <0.05 (**Figure 5D, Supplemental Data 1F**). Gene set enrichment analysis was notable for downregulation of activity-dependent immediate early gene NPAS4-mediated transcription, and downregulation of voltage-gated potassium channels when *FOXP2* levels are low (**Figure 5F, Supplemental Data 1G**). Downregulation of ‘voltage-gated potassium channels’ was a top, common GSEA-pathway between the human postmortem and mouse *Foxp2* KD datasets, consistent with the observed electrophysiological decrease in potassium channel dynamics and increased excitability of the ITCs.

### Validation of transcriptional targets downstream of Foxp2 signaling by qPCR and in situ hybridization

To ensure replicability from our bulk RNAseq analysis, we performed further validation of select transcription targets downstream of Foxp2 signaling. Starting with quantification of normalized counts of three downregulated (*Foxp2, Pbx3, Tac2*) and three upregulated (*Slc12a3, Mesp2, Amh*) genes after *Foxp2* KD observed by bulk RNAseq, (**Figure 6A**) (Unpaired Welch’s t-test, *Foxp2* p<0.0001, *Pbx3* p<0.0001, *Tac2* p<0.0005, *Slc12a3* p<0.0001, *Mesp2* p<0.0001, *Amh* p<0.0001), we performed qPCR on the same RNA as was used for the bulk RNAseq analysis, obtained in a separate aliquot after initial RNA extraction. qPCR replicated up/down regulation for all of the above transcripts, relative to GAPDH expression (Unpaired Welch’s t-test, *Foxp2* p<0.0001, *Pbx3* p=0.0027, *Tac2* p<0.0003, *Slc12a3* p<0.0001, *Mesp2* p<0.0001, *Amh* p<0.0001) (**Figure 6B**). We next validated two separate targets with RNAscope *in situ* hybridization, focusing on one gene with ubiquitous expression in the amygdala (*Oprk1*, encoding the kappa opioid receptor, expressed in the ITCs, basolateral and central amygdala) and one with known differential expression in subnuclei of the amygdala (*Gal*, encoding neuropeptide galanin, expressed in a subset of cells in the central amygdala). We found that *Oprk1* was decreased in downstream targets of the mITCs, including the CeA (unpaired t-test, p=0.002) and basomedial amygdala (unpaired t-test, p=0.001), without a change in the BLA, which acts upstream of ITC signaling during fear learning. *Gal* expression is localized to a subset of central amygdala cells and was significantly decreased after *Foxp2* KD (unpaired t-test, p=0.007), which is consistent with the RNA-seq finding (**Figure 6C**).

**Figure 6.**
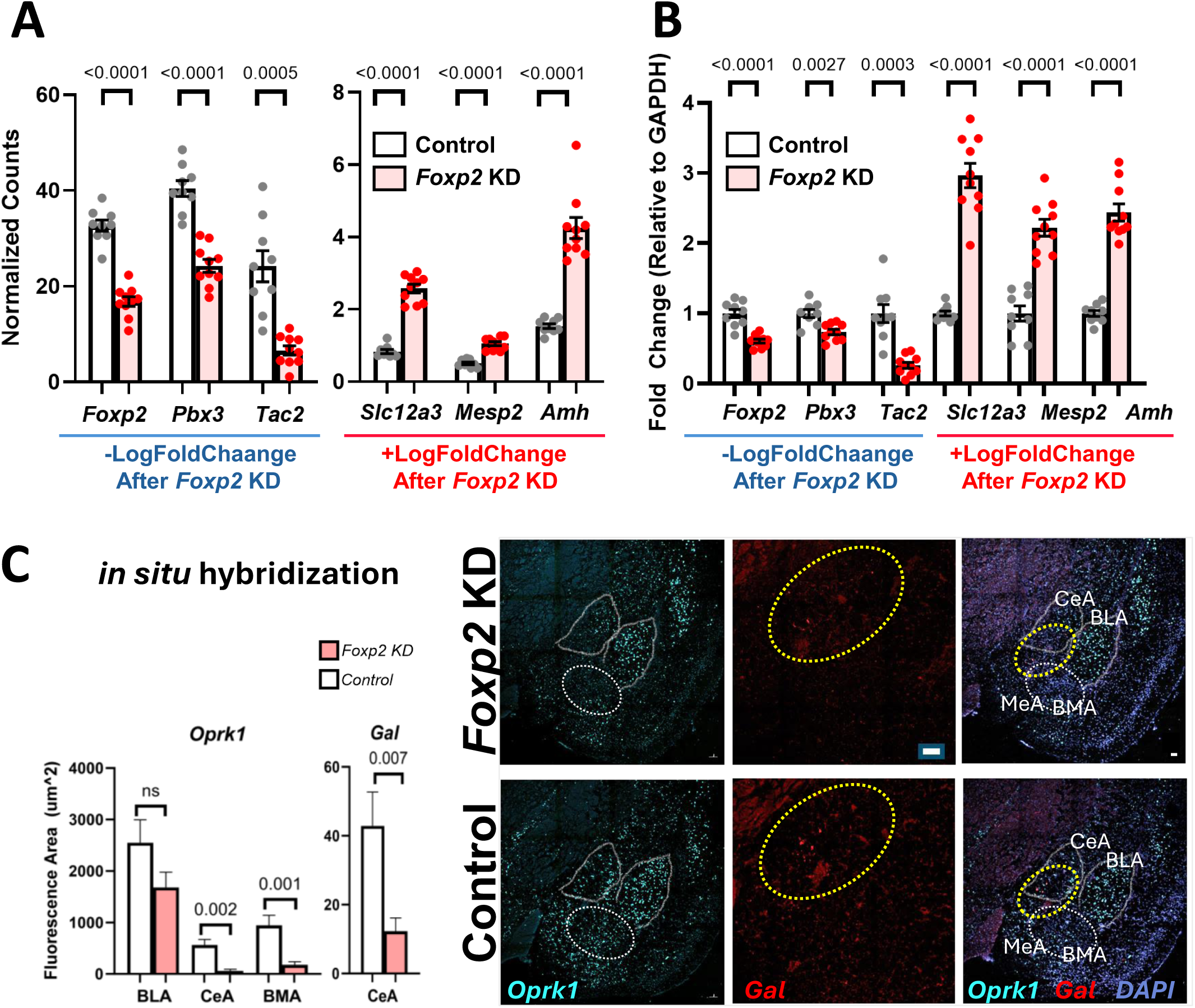
RT-PCR and *in situ* validation of targets downstream of *Foxp2* KD in the amygdala. (A) Bulk RNAseq expression levels of select differentially expressed genes between Foxp2 KD and control groups with normalized gene expression values. Each dot represents an individual mouse. (B) qPCR confirms downregulation of *Foxp2*, *Pbx3, Tac2* and upregulation of *Slc12a3, Mesp2,* and *Amh* following *Foxp2* KD; unpaired Welch’s t-test, n=9 (Control), 10 (Foxp2 KD). (C) RNAscope *in situ* hybridization confirms downregulation of *Oprk1* and *Gal.* BLA (basolateral amygdala), CeA (Central Amygdala), BMA (basomedial amygdala). Unpaired t-test, n=14 amygdala (control), 12 (Foxp2 KD).

Taken together, these results validate the observed up- and down-regulation of mRNA transcription seen in bulk RNA-seq analysis and suggest that widespread transcriptional changes in downstream circuits with ITC projections are mediated by the Foxp2 transcription factor.

### Foxp2 KD decreases signaling through the canonical Wnt pathway

In the brain, transcription of Foxp2 is activated by Lef1, a transcription factor in the canonical WNT/β-catenin signaling, and its downstream effectors, including Pax6, and Pou3Ff2.^39,101–103^ Canonical Wnt signaling, also called Wnt/β-catenin signaling, is initiated when an extracellular Wnt agonist binds to a transmembrane receptor, initiating a cascade that inhibits GSK3β and allows for nuclear translocation of β-catenin, which binds to TCF/LEF transcription factors and activates downstream transcription of Wnt target genes. The alkali metal, lithium, also used as a mood- or emotion-stabilizer in psychiatric disorders (most commonly in bipolar disorder), acts as an intracellular Wnt agonist by binding to GSK3β and inactivating it, thus stimulating nuclear β-catenin signaling^104^ (**Figure 7A**). Foxp2 can also directly interact with β-catenin and thus regulate downstream transcription through the canonical Wnt pathway.^105^ Emerging evidence also suggests that the Wnt/β-catenin pathway directly regulates expression of multiple ion channels in the brain, and potassium channel activity can, in turn, regulate β-catenin levels.^106,107^

**Figure 7.**
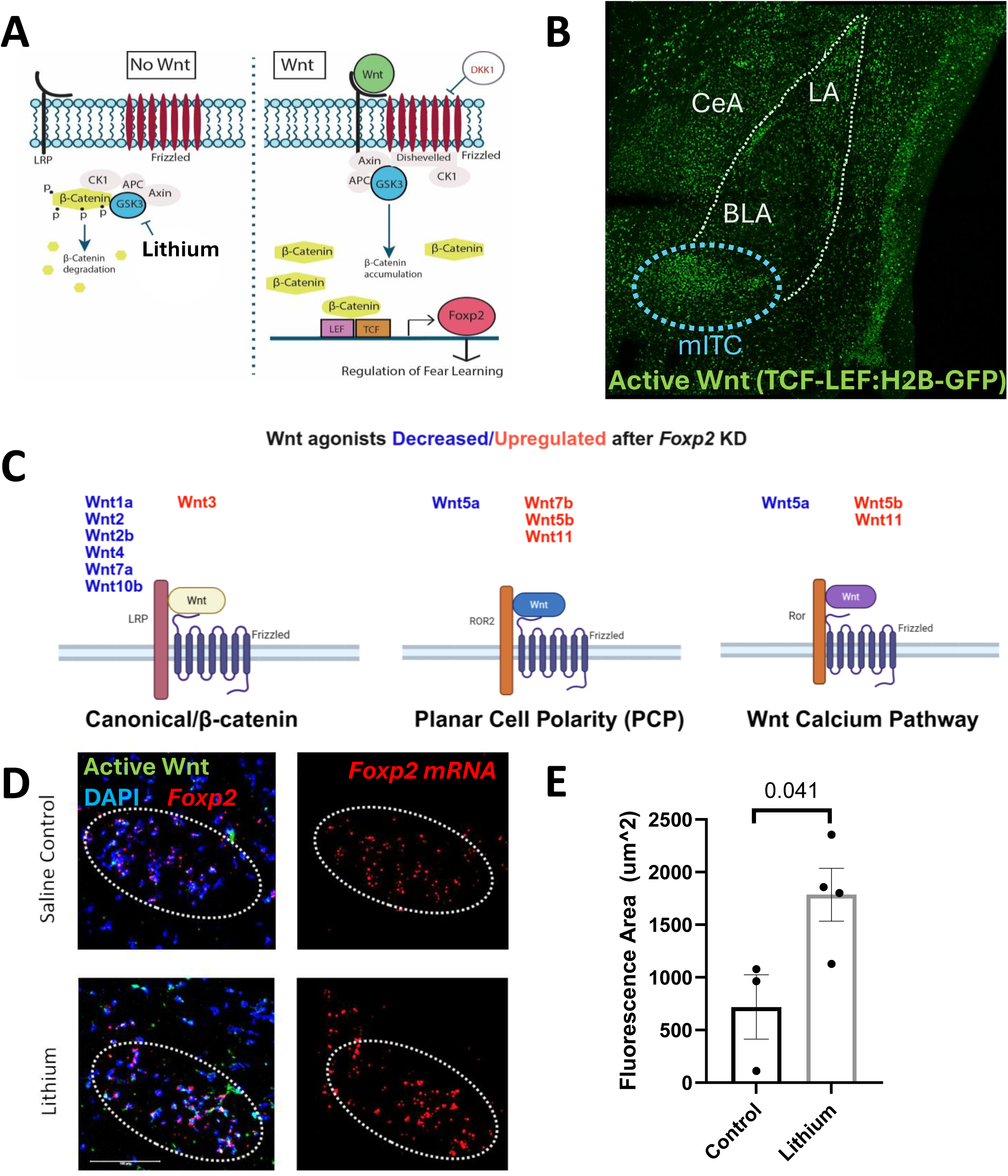
Transcriptional interaction of canonical Wnt and Foxp2 pathways. (A) Hypothesized pathway by which canonical Wnt/β-catenin pathway is regulating *Foxp2* transcription and downstream fear learning. Lithium, an intracellular Wnt agonist, inhibits GSK3β phosphorylation of β-catenin, allowing β-catenin cytoplasmic accumulation, translocation to the nucleus, and stimulation of TCF-LEF transcriptions of Wnt effectors. (B) GFP expression downstream of TCF-LEF transcription factors demonstrating active Wnt signaling in the mITC of the mouse amygdala (blue outline). (C) Extracellular effectors of canonical/b-catenin Wnt signaling are relatively decreased after *Foxp2* KD. (D,E) Wnt agonism with Lithium increases *Foxp2* mRNA levels in the mITC. Unpaired t-test, male and female mice combined. mITC, main Intercalated Cell cluster; BLA, basolateral amygdala; LA, lateral amygdala; CeA, central amygdala.

We previously showed that dynamic signaling from the Wnt/β-catenin pathway is required for fear memory consolidation in mice.^41,76^ Here, we also find that canonical Wnt signaling is expressed in the ITCs in a transgenic fluorescent reported mouse line downstream of TCF/LEF transcription factor (**Figure 7B**). To test for a possible transcriptional interaction between canonical Wnt and Foxp2 pathways in the ITCs during fear learning, we first examined transcriptional regulation downstream of Wnt effectors after *Foxp2* KD (**Figure 7C).** Of the Wnt pathway components from the PANTHER^108^ Wnt signaling pathway dataset (n=254 genes), 136 genes reached FDR significance in our experiment after *Foxp2* KD, suggesting significant enrichment of the Wnt pathway downstream of Foxp2 (*χ*^2^_(1, 254)_ =11.9795, p=0.0005). Of the Wnt pathway components reaching adjusted p-value < 0.05 in the *Foxp2* KD dataset, we found that extracellular effectors of the canonical Wnt/β-catenin signaling pathway were relatively decreased, without a similar decrease in the effectors of the Wnt PCP and Calcium pathways, consistent with our prior findings suggesting that dynamic regulation of the Wnt/β-catenin pathway is required for fear memory consolidation in mice^76^ (**Figure 7C**).

We previously found that lithium increases freezing during extinction of auditory fear conditioning in mice.^76^ We hypothesized that this effect of lithium may be mediated by Wnt signaling regulating downstream transcription of *Foxp2* in the amygdala ITCs, and thus mediating subsequent fear responses. To test this hypothesis, we stimulated Wnt signaling with lithium chloride. We found that lithium (LiCl, I.P., 100 mg/kg administered 30 mins prior to fear learning) increases *Foxp2* mRNA levels, further suggesting that *Foxp2* mRNA is regulated by canonical Wnt signaling in the amygdala (**Figure 7D,E**). Taken together, these results suggest a model in which activated canonical Wnt signaling regulates *Foxp2* transcription in the ITCs, decreasing ITC excitability and disinhibiting its downstream effectors, corresponding to increased freezing after fear memory consolidation. This finding is consistent with a model in which a decrease in *Foxp2* signaling, as we performed with *Foxp2* KD, is associated with decreased freezing levels and reduced fear expression.

### Transcriptional Correlation with Genes Predictive of PTSD

We examined the overlap between transcriptional changes downstream of *Foxp2* from our analysis of the mouse amygdala and genes hypothesized to be causative in PTSD from large-scale human GWAS (Tier 1 genes, defined as being putatively PTSD causal, and Tier 2 genes, defined as genes reaching GWAS significance, but less likely to be putative causal when compared with Tier 1 genes).^8^ Of the 129 putative Tier 1 and Tier 2 GWAS-significant PTSD causal genes identified by Nievergelt et al, 108 genes had mouse orthologues that were detected in our list. Of these, 43 genes reached p-adjusted value of <0.01 after *Foxp2* KD, pointing to a substantial enrichment of putative PTSD-causative genes that were differentially expressed in our amygdala dataset following *Foxp2* KD (*χ*^2^ _(1, N=108)_ = 4.954, p= 0.026) (**Figure 8A,C**).

**Figure 8.**
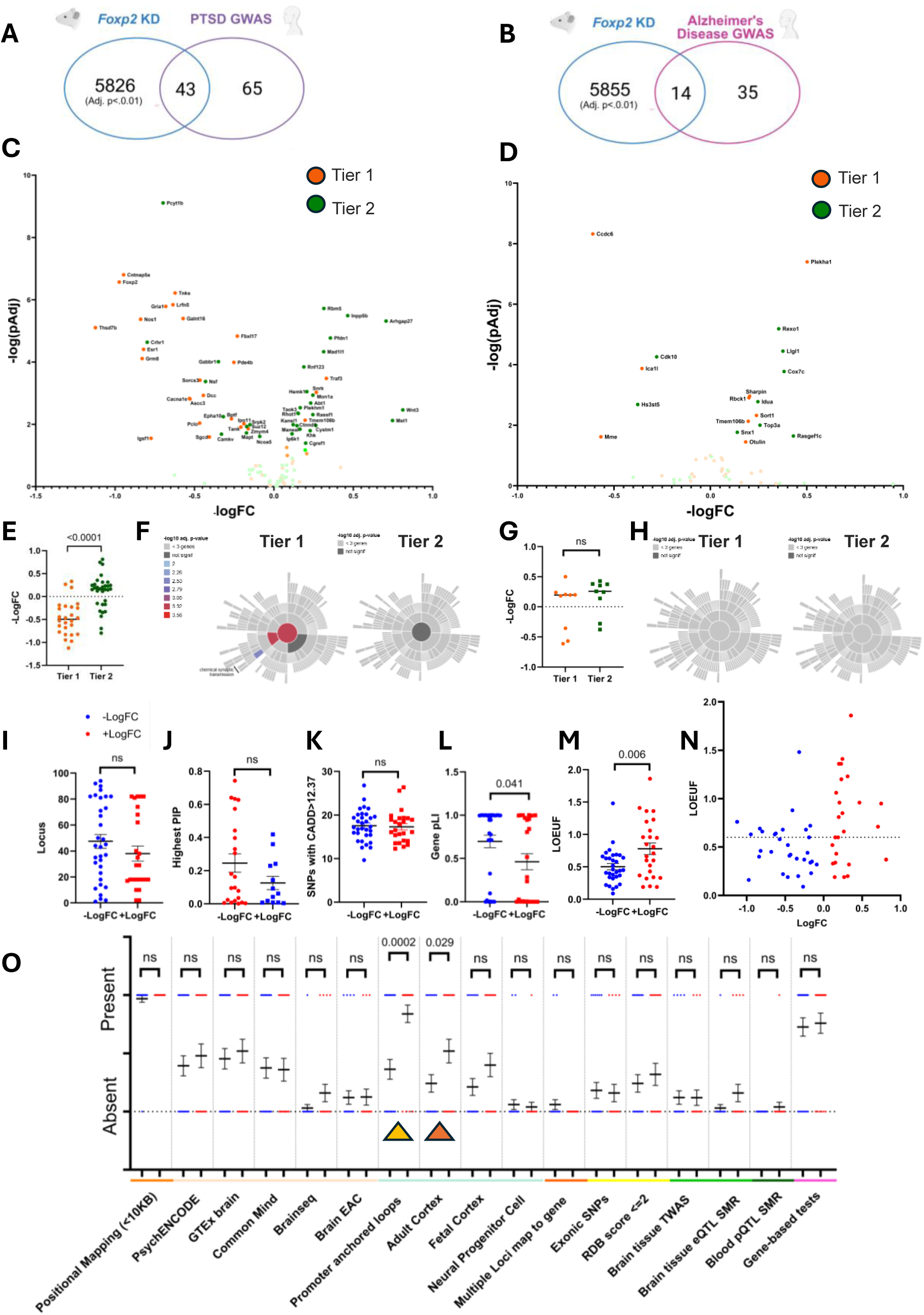
*Foxp2* KD DEGs are enriched for putatively PTSD-causal genes. *Foxp2* KD dataset in the mouse amygdala was compared to putative causal genes from PTSD (A) and Alzheimer’s Disease (AD), (B) GWAS. Number of putative causal genes are compared with up and downregulated DEGs at adjusted significance level of p<0.01, and show significant enrichment in PTSD, but not AD, datasets. Volcano plot highlighting *Foxp2* KD DEGs that were also identified as being putative high risk (Tier 1) and risk (Tier 2) genes for PTSD (C) and AD (D), with genes reaching adj p<0.05 labeled in darker colors. (E) Mouse orthologue Tier 1 genes from PTSD GWAS are significantly downregulated after *Foxp2* KD and enriched in synaptic pathways (F), while Tier 2 genes are relatively upregulated and show no synaptic enrichment. Mouse orthologue Tier 1 and Tier 2 genes from the AD GWAS show no difference in up or downregulation (G), and no synaptic enrichment (H) after *Foxp2* KD. Individual components of tiered gene ranking from the PTSD GWAS are shown in (I-L, O), with *Foxp2* KD mouse orthologues of Tier 1 and Tier 2 PTSD genes by individual tiered gene ranking components, as defined in Nievergelt et al.^8^ We find no difference in the locus location (I). Highest Posterior Importance Probability (PIP) (J), and Combined annotation dependent depletion (CADD) scores for GWAS prioritized genes among mouse orthologues downregulated or upregulated after *Foxp2* KD (K). Tiered gene pLI (probability of Loss of Impact)^110^, was increased after *Foxp2* KD. (L) GWAS prioritized genes downregulated after *Foxp2* KD have overall lower LOEUF scores (M) with a trend toward enrichment of genes with LOEUF scores below 0.6 (N). Additional evidence categories for tiered ranking for human PTSD GWAS targets shown as presence/absence of the listed domains in the downregulated (blue) or upregulated (red) mouse orthologues after *Foxp2* KD. Significant differences were noted in the presence of promoter anchored loops (highlighted with yellow arrowhead) and transcripts in the adult cortex (highlighted with red arrowhead). (O). Abbreviations: RDB, Regulome database; TWAS, transcriptome-wide association study; eQTL, expression quantitative trait locus; pQTL, protein quantitative trait locus; SMR, summary Mandelian randomization. All statistics represent unpaired t-tests, unless otherwise indicated.

To ensure that the observed enrichment was not due to general transcriptional changes that could be observed in CNS pathology, such as transcriptional activation of inflammatory pathways, we compared our *Foxp2* KD DEG dataset to GWAS of Alzheimer’s Disease (AD) with similar Tier 1/Tier 2 putative disease risk stratification.^82,109^ In contrast to the enrichment observed in the *Foxp2* KD dataset with putative causal genes identified in the PTSD GWAS, we observed no enrichment in genes from the AD dataset. Of the 55 putative Tier 1 and Tier 2 AD causal genes identified by Bellenguez et al,^82^ 49 genes had mouse orthologues that were detected in our list. Of these, 14 genes were identified as reaching p-adjusted significance of <0.01 after *Foxp2* KD, showing no enrichment of putative AD-causative genes in our dataset above that which would be expected by chance (*χ*^2^ _(1, N=49)_ = 0.048, p= 0.8273). (**Figure 8B, D**)

Of note, genes identified as Tier 1 from the PTSD GWAS were more likely to be downregulated, while genes identified as Tier 2 were more likely to be upregulated after *Foxp2* KD (p<0.0001) (**Figure 8E**), whereas no difference was observed in up or downregulation of Tier 1 vs Tier 2 genes from the AD GWAS after *Foxp2* KD (**Figure 8G**). This result further emphasizes the specificity in enrichment of targets implicated in PTSD downstream of Foxp2 signaling in the amygdala.

Pathway enrichment analysis of PTSD Tier 1 p-adjusted significant *Foxp2* KD genes in SynGO revealed 8/25 genes that mapped to unique SynGO annotated genes (*PCLO, CACNA1E, GRIA1, DCC, GRM8, NOS1, LRFN5, SORCS3*). Relative to other genes expressed in the brain, these genes were overrepresented in biological processes at the synapse (p= 1.26×10^−4^, q= 5.03×10^−4^), synaptic signaling (p= 2.87×10^−4^, q= 5.75×10^−4^), and chemical synaptic transmission (p= 2.24×10^−3^, q= 2.99×10^−3^). By contrast, 5 of the 33 PTSD Tier 2 / p-adjusted significant *Foxp2* KD genes mapped to unique SynGO annotated genes (*CAMKV, CRHR1, GABBR1, NSF, SRPK2*); however, there were no significantly enriched genes at the 1% FDR level (testing term with at least three matching input genes) for cellular component or biological processes (**Figure 8F**). We also found no synaptic enrichment for mouse orthologs of tiered AD genes in our *Foxp2* KD dataset (**Figure 8H**).

To further explore the finding that top tier PTSD putative causal genes are generally downregulated after *Foxp2* KD, we separated individual components of tiered gene ranking from the PTSD GWAS to probe their relative contribution to enrichment of up or downregulated targets after *Foxp2* KD.^8^ We found no difference in locus location of the human ortholog, (**Figure 8I**) in the highest Posterior Importance Probability (PIP), a measure that estimates the probability that a given gene variant may be causally involved in PTSD based on fine-mapping, (**Figure 8J**) or in the Combined Annotation Dependent Depletion (CADD) scores for GWAS-prioritized genes among mouse orthologues downregulated or upregulated after *Foxp2* KD (**Figure 8K**). Tiered gene probability of Loss of Impact (pLI)^110^ a measure of population sensitivity to heterozygous loss of function mutations, was increased after *Foxp2* KD, however, was not significant at pLI>0.9, albeit with a moderate association suggested by Cramer’s V effect size (*χ*²(1) = 2.65, p = 0.103, Cramér’s V = 0.21) (**Figure 8L**). To follow up on this finding, we analyzed LOEUF scores (Loss-of-function observed/Expected upper bound fraction)^83^ from gnomADv2.1+ of the prioritized PTSD GWAS orthologues after *Foxp2* KD. We found that GWAS prioritized genes downregulated after *Foxp2* KD have overall lower LOEUF scores (**Figure 8M**), with a trend toward enrichment of genes with LOEUF scores below 0.6 (*χ*²(1, 55) = 3.4774, p = 0.06) (**Figure 8M, N**). This result suggests that mouse orthologs of top PTSD tier genes that are transcriptionally decreased after *Foxp2* KD may be more intolerant to loss-of-function mutations, which is consistent with the evolutionary conservation of Foxp2 and points to an enrichment of functionally critical genes downstream of Foxp2 transcriptional regulation.

*Foxp2* KD mouse orthologues of tiered PTSD GWAS genes with additional evidence categories from the tiered ranking, including components of positional mapping, eQTL mapping, and chromatin interaction mapping, are shown as presence/absence of the listed domains in **Figure 8O**. We found that *Foxp2* KD decreases transcription of mouse orthologues of PTSD GWAS tiered genes with fewer promoter chromatin anchored loops, as well as genes that are less likely to be expressed in the adult cortex.

Overall, these results suggest that downregulated genes enriched for human orthologs of Tier 1 PTSD GWAS risk genes reflect subcortical FOXP2/Foxp2 signaling within the amygdala, and that FOXP2/Foxp2 may directly bind regulatory elements of top PTSD risk genes, rather than acting through distal enhancer-promoter looping.

## Discussion

Prior work has implicated the role of FOXP2 in a number of psychiatric symptoms and illnesses, including schizophrenia, ADHD, bipolar disorder, and PTSD.^8,12,111^ Our work demonstrates a vital role for the Foxp2 transcription factor - the top gene genetically associated with PTSD to date - in mediating fear memory expression through transcriptional regulation in amygdala ITCs. In mammals, the amygdala serves as a hub for formation, consolidation, and expression of emotional memories, including regulation of fear learning memory, with FOXP2/Foxp2 highly expressed in amygdala ITCs that surround and regulate the basolateral amygdala. To date, multiple studies have used *Foxp2* as a marker for circuit studies of ITCs, which act as mediators between the basolateral amygdala and subsequent behavioral output, mediated by the central amygdala. However, the role of *Foxp2* and its downstream molecular pathways in the amygdala has not been known. In the present study, we show that a decrease in *Foxp2* signaling leads to increased excitability of the ITCs, which is consistent with increased inhibition of the central amygdala and its downstream targets, resulting in a decrease in observed behavioral response to fear. We further show that *Foxp2* acts as a hub gene for transcriptional regulation of multiple molecular pathways previously identified in animal models of fear learning and human genomic studies of PTSD (**Figure 9**).

**Figure 9.**
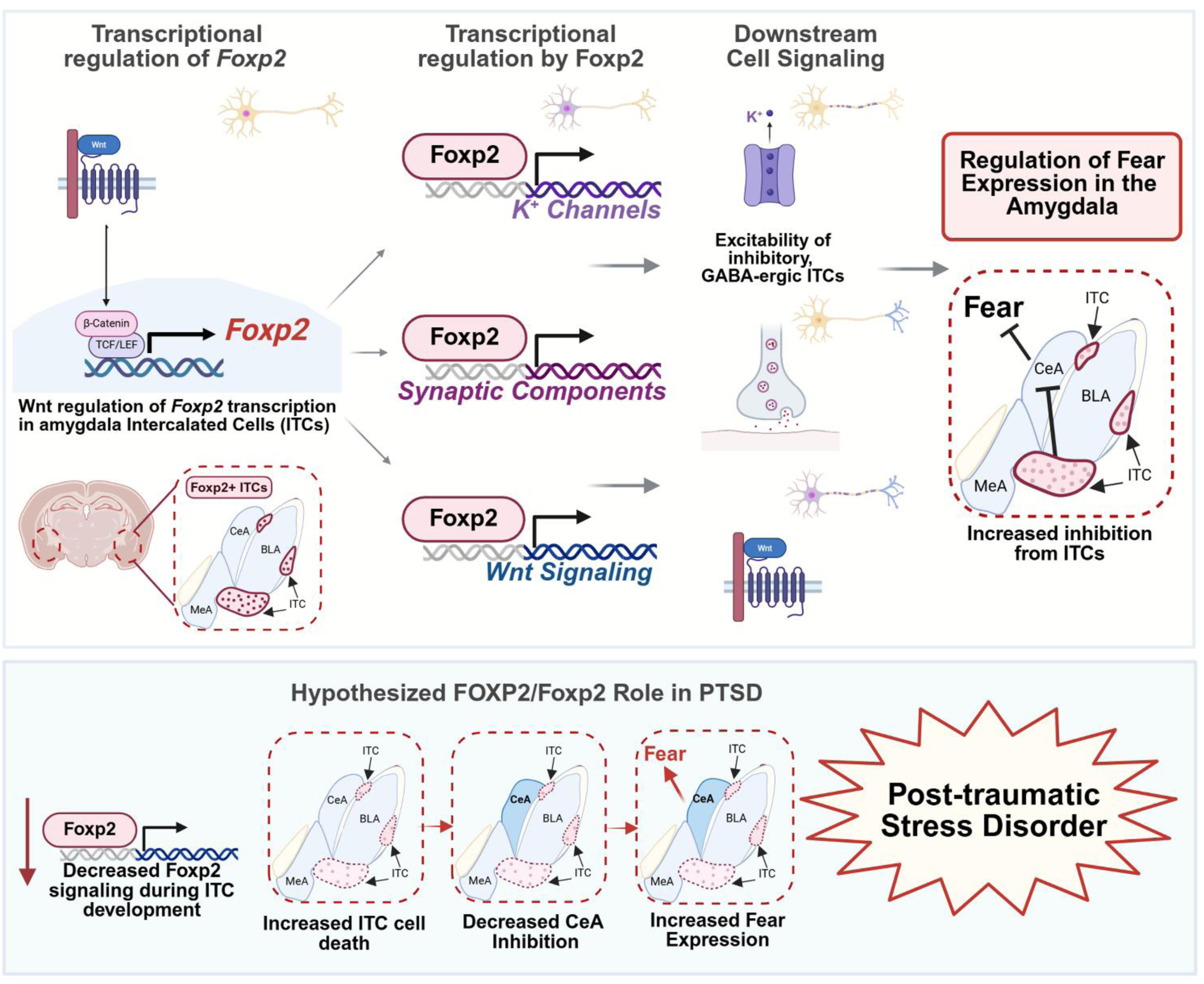
Transcriptional regulation by *Foxp2* controls ITC excitability and downstream expression of fear learning. Chronic decreased expression of FOXP2/Foxp2 is associated with increased ITC cell death during development, likely contributing to decreased CeA inhibition and increased baseline expression of fear, leading to development of PTSD in the setting of a stress trigger. Other possibilities include regulatory SNPs that lead to dynamic alterations in *Foxp2* expression in adults in the presence of trauma, leading to dysregulated amygdala function increasing risk for trauma-related disorders.

Curiously, transcriptomic data from large-scale human studies suggest that intronic *FOXP2* SNPs associated with PTSD are also associated with a decrease in *FOXP2* transcription. However, we find that that temporary *Foxp2* knockdown in the amygdala was associated with a decrease in expression of fear to an acute stressor. There are several likely explanations that reconcile these observations. First, in humans *FOXP2* levels would be decreased throughout the lifespan of the individual, including during critical windows of amygdala and fear circuitry formation. Foxp2 has a well-established role in development, including in regulation of neuronal maturation and interaction with Wnt signaling, which also plays a vital role in regulation of axon guidance and synapse formation pathways.^112–117^ A reduction in *Foxp2* signaling during amygdala development in mice leads to ITC cell death and lack of ITC circuit formation.^84^ Thus, a lifelong reduction in FOXP2 signaling would likely be associated with underdeveloped ITC circuits and a hyperexcitable amygdala. This hypothesis is consistent with our observed decrease in inhibitory neurons, which is associated with decreased *FOXP2* levels in homozygous carriers of PTSD risk-associated *FOXP2* SNPs. To remove the contribution of underdeveloped amygdala circuit in the context of developmental *Foxp2* reduction on fear learning, we focused on characterization of an acute, region-specific reduction in a previously-formed circuit.

Another possible explanation for the observed difference in chronic vs acute FOXP2/Foxp2 reduction is compensatory sensitization or compensation from chronic reduction of *FOXP2*. A similar phenomenon was observed in the BNST with a reduction of CRFR1 receptor density after overexpression of the corticotropin releasing factor (CRF).^118^ Broadly, genetic compensation is defined as changes in RNA or protein levels of genes that functionally compensate for the substantial decrease or loss of function of the targeted gene. A subject of active investigation, genetic compensation has been described in multiple animal models, including mice, and has in some situations resulted in lack of observed phenotype after complete loss of function.^119^ More recently, genetic compensation from Foxp1 following loss of Foxp2, has been demonstrated in the striatum.^120^ To further decrease the potential for genetic compensation after Foxp2 knockout, we chose a conditional knockdown approach. Tissue-specific transcriptional regulation depends on co-expression of paralogues Foxp1, Foxp2, and Foxp4, which homo and heterodimerize prior to DNA binding.^121^ Only Foxp2 and Foxp1 are expressed into adulthood and, while strongly expressed in the striatum, Foxp1 is not expressed in the ITCs, suggesting that any compensation/interaction of Foxp2 in the amygdala has yet to be defined. If reduction of Foxp2 levels in the ITCs is not compensated by a paralogue binding such as Foxp1, then amygdala Foxp2 may be especially sensitive to small disruptions in Foxp2 levels, as would be expected from changes in regulatory SNPs, as observed in the PTSD GWAS.

We focused on the main ITC with an initial hypothesis that Foxp2’s role would be most robustly observed during fear extinction, given the previously characterized role for mITC in extinction and extinction retrieval.^122^ To our surprise, KD of Foxp2 in the amygdala resulted in a significant decrease in fear expression during both fear acquisition and extinction phases. This decrease was correlated with increased physiological excitability of the ITCs, suggesting increased feedforward inhibition to the CeA, which should be tested in future studies.

Increased inhibition of the CeA, and specifically the medial central amygdala, to which the mITCs send direct projections, would decrease output to the brainstem and the hypothalamus, dampening the freezing response to a conditioned stimulus. Our viral KD construct was driven by a ubiquitous neuronal promoter, and KD specificity is driven primarily by limited localization of Foxp2 expression in the ITCs of the amygdala, rather than in the BLA or the CeA. Although we targeted the mITC cluster, recent evidence suggests interconnectivity of ITCs.^80,122,123^ Thus, we cannot exclude that Foxp2 KD targeting one cluster did not have downstream transcriptional responses in other ITCs.

The observed transcriptional regulation by Foxp2 also includes regions of the amygdala outside of the ITCs, which would have been included in the brain punches, such as the BLA, CeA, and small parts of the medial amygdala. These regions may mediate key pathways previously identified in fear learning, including *Tac2, CRH,* vasopressin, multiple components regulating synapse function, including *TNKS* and *PCLO*, as well as GABA/GIRK signaling previously identified as being affected by *Foxp2* KD.^29^ When compared with recently prioritized genes in a PTSD GWAS meta-analysis, downregulated transcripts downstream of *Foxp2* in the amygdala with the highest significance appear to be enriched for genes identified as putatively PTSD causal in the GWAS analysis (Tier 1 genes), while Tier 2 genes (also reaching GWAS significance but less likely putative causal genes) are relatively enriched among genes upregulated after *Foxp2* KD. This separation appears to reflect subcortical localization of Tier 1 gene signaling, which may reflect the location of both FOXP2-expressing ITCs, and its downstream projections in the amygdala.^64,124^ Further, decreased expression of tiered genes without promoter-anchored chromatin loops after *Foxp2* KD suggests that the Foxp2 transcription factor may be directly binding proximal, cis-regulatory elements of top tier PTSD risk genes as a central regulator of some of these genetically associated pathways. Together, these results support a hypothesis in which top PTSD risk gene *FOXP2* acts as a transcriptional hub targeting a network of genes that are, themselves, independently enriched for PTSD genetic risk. This model suggests that both transcriptional regulation and genetic variation converge on the same molecular pathways to amplify PTSD susceptibility.

In conclusion, our results suggest that Foxp2 acts as a hub transcription factor to regulate expression of multiple fear-learning related pathways in the amygdala. Follow up mechanistic studies will be needed to define pathways by which Foxp2 regulates fear expression, which will likely elucidate novel treatment targets for PTSD and other forms of anxiety, trauma- and stress-related pathology.

## Acknowledgements

We would like to thank Khalil Threadgill and Emily Newman for their assistance with stereotaxic surgeries, Jakob Hartmann for assistance with viral construct design, Edward Meloni for assistance with the acoustic startle test, Murray Stein for feedback and service to the PGC-PTSD working group, as well as Kwang-Soo Kim, Gary Kaplan, Ann Rasmusson, Yan Li, Beibei Peng, and Vadim Bolshakov for helpful discussions relating to data interpretation. We also thank the late Cristina Berciu for her help with confocal imaging and the late Elena Chartoff for her insights and support of this project.

We acknowledge the members of the PTSD Working Group of Psychiatric Genomics Consortium Caroline M. Nievergelt, Adam X. Maihofer, Elizabeth G. Atkinson, Chia-Yen Chen, Karmel W. Choi, Jonathan R. I. Coleman, Nikolaos P. Daskalakis, Laramie E. Duncan, Renato Polimanti, Cindy Aaronson, Ananda B. Amstadter, Soren B. Andersen, Ole A. Andreassen, Paul A. Arbisi, Allison E. Ashley-Koch, S. Bryn Austin, Esmina Avdibegoviç, Dragan Babić, Silviu-Alin Bacanu, Dewleen G. Baker, Anthony Batzler, Jean C. Beckham, Sintia Belangero, Corina Benjet, Carisa Bergner, Linda M. Bierer, Joanna M. Biernacka, Laura J. Bierut, Jonathan I. Bisson, Marco P. Boks, Elizabeth A. Bolger, Amber Brandolino, Gerome Breen, Rodrigo Affonseca Bressan, Richard A. Bryant, Angela C. Bustamante, Jonas Bybjerg-Grauholm, Marie BækvadHansen, Anders D. Børglum, Sigrid Børte, Leah Cahn, Joseph R. Calabrese, Jose Miguel Caldas-de-Almeida, Chris Chatzinakos, Sheraz Cheema, Sean A. P. Clouston, Lucía Colodro-Conde, Brandon J. Coombes, Carlos S. Cruz-Fuentes, Anders M. Dale, Shareefa Dalvie, Lea K. Davis, Jürgen Deckert, Douglas L. Delahanty, Michelle F. Dennis, Frank Desarnaud, Christopher P. DiPietro, Seth G. Disner, Anna R. Docherty, Katharina Domschke, Grete Dyb, Alma Džubur Kulenović, Howard J. Edenberg, Alexandra Evans, Chiara Fabbri, Negar Fani, Lindsay A. Farrer, Adriana Feder, Norah C. Feeny, Janine D. Flory, David Forbes, Carol E. Franz, Sandro Galea, Melanie E. Garrett, Bizu Gelaye, Joel Gelernter, Elbert Geuze, Charles F. Gillespie, Slavina B. Goleva, Scott D. Gordon, Aferdita Goçi, Lana Ruvolo Grasser, Camila Guindalini, Magali Haas, Saskia Hagenaars, Michael A. Hauser, Andrew C. Heath, Sian M. J. Hemmings, Victor Hesselbrock, Ian B. Hickie, Kelleigh Hogan, David Michael Hougaard, Hailiang Huang, Laura M. Huckins, Kristian Hveem, Miro Jakovljević, Arash Javanbakht, Gregory D. Jenkins, Jessica Johnson, Ian Jones, Tanja Jovanovic, Karen-Inge Karstoft, Milissa L. Kaufman, James L. Kennedy, Ronald C. Kessler, Alaptagin Khan, Nathan A. Kimbrel, Anthony P. King, Nastassja Koen, Roman Kotov, Henry R. Kranzler, Kristi Krebs, William S. Kremen, Pei-Fen Kuan, Bruce R. Lawford, Lauren A. M. Lebois, Kelli Lehto, Daniel F. Levey, Catrin Lewis, Israel Liberzon, Sarah D. Linnstaedt, Mark W. Logue, Adriana Lori, Yi Lu, Benjamin J. Luft, Michelle K. Lupton, Jurjen J. Luykx, Iouri Makotkine, Jessica L. Maples-Keller, Shelby Marchese, Charles Marmar, Nicholas G. Martin, Gabriela A. Martínez-Levy, Kerrie McAloney, Alexander McFarlane, Katie A. McLaughlin, Samuel A. McLean, Sarah E. Medland, Divya Mehta, Jacquelyn Meyers, Vasiliki Michopoulos, Elizabeth A. Mikita, Lili Milani, William Milberg, Mark W. Miller, Rajendra A. Morey, Charles Phillip Morris, Ole Mors, Preben Bo Mortensen, Mary S. Mufford, Elliot C. Nelson, Merete Nordentoft, Sonya B. Norman, Nicole R. Nugent, Meaghan O’Donnell, Holly K. Orcutt, Pedro M. Pan, Matthew S. Panizzon, Gita A. Pathak, Edward S. Peters, Alan L. Peterson, Matthew Peverill, Robert H. Pietrzak, Melissa A. Polusny, Bernice Porjesz, Abigail Powers, Xue-Jun Qin, Andrew Ratanatharathorn, Victoria B. Risbrough, Andrea L. Roberts, Alex O. Rothbaum, Barbara O. Rothbaum, Peter Roy-Byrne, Kenneth J. Ruggiero, Ariane Rung, Heiko Runz, Bart P. F. Rutten, Stacey Saenz de Viteri, Giovanni Abrahão Salum, Laura Sampson, Sixto E. Sanchez, Marcos Santoro, Carina Seah, Soraya Seedat, Julia S. Seng, Andrey Shabalin, Christina M. Sheerin, Derrick Silove, Alicia K. Smith, Jordan W. Smoller, Scott R. Sponheim, Dan J. Stein, Synne Stensland, Jennifer S. Stevens, Jennifer A. Sumner, Martin H. Teicher, Wesley K. Thompson, Arun K. Tiwari, Edward Trapido, Monica Uddin, Robert J. Ursano, Unnur Valdimarsdóttir, Miranda Van Hooff, Eric Vermetten, Christiaan H. Vinkers, Joanne Voisey, Yunpeng Wang, Zhewu Wang, Monika Waszczuk, Heike Weber, Frank R. Wendt, Thomas Werge, Michelle A. Williams, Douglas E. Williamson, Bendik S. Winsvold, Sherry Winternitz, Christiane Wolf, Erika J. Wolf, Yan Xia, Ying Xiong, Rachel Yehuda, Keith A. Young, Ross McD Young, Clement C. Zai, Gwyneth C. Zai, Mark Zervas, Hongyu Zhao, Lori A. Zoellner, John-Anker Zwart, Terri deRoon-Cassini, Sanne J. H. van Rooij, Leigh L. van den Heuvel, AURORA Study, Estonian Biobank Research Team, FinnGen Investigators, HUNT All-In Psychiatry, Murray B. Stein, Kerry J. Ressler and Karestan C. Koenen

We acknowledge the member of the PsychENCODE PTSD BrainOmics Project: Dhivya Arasappan, Sabina Berretta, Rahul A. Bharadwaj, Frances A. Champagne, Leonardo Collado-Torres, Christos Chatzinakos, Nikolaos P. Daskalakis, Chris P. DiPietro, Duc M. Duong, Amy Deep-Soboslay, Nick Eagles, Louise Huuki, Thomas Hyde, Artemis Iatrou, Aarti Jajoo, Joel E. Kleinman, Charles B. Nemeroff, Geo Pertea, Kerry J. Ressler, Deanna Ross, Nicholas T. Seyfried, Joo Heon Shin, Clara Snijders, Ran Tao, Daniel R. Weinberger, Stefan Wuchty, Dennis Wylie

## Author contributions

Conceptualization: OP, KJR; Data curation: OP, LTS, MMM, CK; Formal analysis: OP, LTS, MMM, CK, VB, RS, ZB, MM, AL; Funding acquisition: OP, KJR, NPD, LTS; Investigation: OP, LTS, CK, SK, EC, JZ, PM; Methodology: OP, LTS, NPD, KJR; Project administration: KJR, OP; Resources: KJR, NPD, QF-B, JS; Supervision: KJR, NPD, WAC; Writing – original draft: OP, LTS, MMM; Writing – review & editing: all

